# Augmenting Neutralization breadth against Diverse HIV-1 by increasing the Ab-Ag interface on V2

**DOI:** 10.1101/2021.08.07.455519

**Authors:** Nan Gao, Yanxin Gai, Lina Meng, Chu Wang, Wei Wang, Xiaojun Li, Tiejun Gu, Mark K. Louder, Nicole A. Doria-Rose, Kevin Wiehe, Alexandra F. Nazzari, Adam S. Olia, Jason Gorman, Reda Rawi, Wenmin Wu, Clayton Smith, Htet Khant, Natalia de Val, Bin Yu, Junhong Luo, Haitao Niu, Yaroslav Tsybovsky, Huaxin Liao, Thomas B. Kepler, Peter D. Kwong, John R. Mascola, Chuan Qin, Tongqing Zhou, Xianghui Yu, Feng Gao

**Affiliations:** National Engineering Laboratory for AIDS Vaccine, School of Life Sciences, Jilin University, Changchun, Jilin Province 130012, China; Institute of Laboratory Animal Science, Chinese Academy of Medical Sciences, Beijing 100021, China; Comparative Medicine Center, Peking Union Medical College, Beijing 100021, China; Department of Medicine, Duke University School of Medicine, Durham, North Carolina 27710, USA; Vaccine Research Center, National Institute of Allergy and Infectious Diseases, National Institutes of Health, Bethesda, MD 20892, USA; Duke University Human Vaccine Institute, Duke University School of Medicine, Durham NC 27710, USA; Cancer Research Technology Program Frederick National Laboratory for Cancer Research, Leidos Biomedical Research Inc., Frederick, MD 21701, USA; School of Medicine, Jinan University, Guangzhou, Guangdong Province 510632, China; Department of Microbiology, Boston University, Boston, Massachusetts 02215, USA; Key Laboratory for Molecular Enzymology and Engineering, the Ministry of Education, School of Life Sciences, Jilin University, Changchun, Jilin Province 130012, China

**Keywords:** HIV-1, SHIV, broadly neutralizing antibodies, maturation, non-human primate

## Abstract

Understanding maturation pathways of broadly neutralizing antibodies (bnAbs) against HIV-1 in non-human primates can be highly informative for HIV-1 vaccine development. We now obtained a lineage of J038 from Chinese rhesus macaques after 7-years of SHIV infection. J038 has short complementary determining loops and neutralizes 54% of global circulating HIV-1 strains. Its binding induces a unique “up” conformation for one of the V2 loops in the trimeric envelope glycoprotein (Env) and is heavily dependent on glycan, which provides nearly half of the binding surface. The unmutated common ancestor of the J038 lineage antibodies binds monomeric gp120 and neutralizes the autologous virus. Continuous maturation enhances neutralization potency and breadth of J038 lineage antibodies via expanding antibody-Env contact areas surrounding the core region contacted by germline-encoded residues. Developmental details and recognition features of J038 lineage antibodies revealed here provide a new pathway for maturation elicitation of V2-targeting bnAbs.

**Highlights:**

• Long-term infected NHPs develop antibodies neutralizing up to 54% of HIV-1 strains
• Antibody J038 binds one V2 loop on HIV-1 Env trimer in a unique “up” position
• UCA of the J038 lineage effectively neutralizes the autologous virus
• J038 lineage antibodies mature through gradually increased contact to glycans

## Introduction

HIV/AIDS has affected 76 million people in the world since the start of the epidemic (UNAIDS, 2021). However, after over 30 years of search, an AIDS vaccine is still illusive. The RV144 phase III trial shows a 31% efficacy (Rerks-Ngarm et al., 2009), but the recent failure of a similar phase III trial in Africa (HVTN 702) shows the extreme challenges in development of an effective AIDS vaccine (Gray et al., 2021). One major challenge is the extraordinarily high level of genetic variation among global circulating HIV-1 strains; up to 30% differences in the envelope glycoprotein (Env) which is the sole target for neutralizing antibodies (nAbs) (Malim and Emerman, 2001).. Thus, broadly neutralizing antibodies (bnAbs) against diverse HIV-1 strains should be required for a vaccine to effectively prevent HIV-1 infection by the global strains (Burton and Mascola, 2015; Haynes et al., 2019; Kwong and Mascola, 2018). Broad neutralization activity can be detected in the sera of ∼20% HIV-1 infected individuals after 2-4 years of infection (Hraber et al., 2014; Landais et al., 2016). Hundreds of bnAbs have been isolated from such individuals and structural analysis demonstrates that six main conserved sites (V2 apex, the CD4 binding site [CD4bs], V3-glycan, the membrane proximal external region [MPER], silence face and the gp120-gp41 interface, including the fusion peptide) on Env are targeted by bnAbs (Kwong and Mascola, 2018; Sok and Burton, 2018). Importantly, infusion of bnAbs in transgenic mice and non-human primates (NHPs) can prevent acquisition of SHIV infection (Gautam et al., 2016; Hessell and Haigwood, 2015; Pegu et al., 2017), indicating that bnAbs can effectively prevent HIV-1 infection if they are elicited by vaccines. The results from the Antibody-Mediated Prevention (AMP) trail shows that VRC01, a CD4bs bnAb, is 75.4% effective to prevent infection of VRC01 sensitive viruses, although overall it does not prevent acquisition of other HIV-1 isolates (Corey et al., 2021). A recent study demonstrates that high titer bnAbs, not T cell immunity or ADCC, is associated with protection of inquisition of SHIV infection (Pauthner et al., 2019). However, such potent and broad nAbs have not be successfully elicited by vaccines in NHPs or humans (Andrabi et al., 2018; Haynes et al., 2019).

Potent neutralization against autologous tier 2 virus has been elicited in animal models by current vaccine strategies (Havenar-Daughton et al., 2016; Sanders et al., 2015; Williams et al., 2017). Recent studies show that some levels of broad neutralization breadth can be achieved. The fusion peptide (FP) vaccine has elicited weak but cross-clade neutralizing activities in mice, guinea pigs and non-human primates (NHPs) (Kong et al., 2019; Xu et al., 2018). A vaccine approach using the N-glycan removal trimers prime and boosting with gradual N-glycan restoration heterologous trimers on liposomes to elite broadly neutralizing responses in rabbits, in which an interface targeted monoclonal bnAb with 87% neutralization breadth was isolated (Dubrovskaya et al., 2019). Other studies show that V3-glycan targeted neutralizing antibodies with limited breadth can be elicited in rhesus macaques (Alam et al., 2017; Han et al., 2019; Sanders et al., 2015). These results show that neutralizing antibodies are likely be induced by vaccine strategies.

NHPs are the only model which can evaluate protection of inquisition of HIV-1 infection (Rahman and Robert-Guroff, 2019). However, little is known about if bnAbs with similar breadth and potency as those in humans can develop in NHPs and whether they developed in a similar manor as in humans (Roark et al., 2021; Wang et al., 2020). A better understanding of these questions will be critical for use of NHPs to evaluate the protection efficacy of HIV-1 Env-based vaccines. In our previous study, we have detected broad neutralization activities against HIV-1 in Chinese rhesus macaques after 6- years of SHIV infection (Gao et al., 2018). Epitope mapping showed that the broad neutralization activity has multiple specificities, targeting V2, V3 and CD4bs that are commonly detected for bnAbs in humans (Gao et al., 2020; Gao et al., 2018). Here we isolated the multiple mAbs from one Chinese rhesus macaque using V2 differential baits. Biological and structural analyses of these mAbs showed that they are broadly neutralizing, uniquely bind to the V2 apex, and have a novel maturation pathway for broad neutralization. These findings will have important implications in understanding the differences in bnAb maturation in humans and NHPs and how to best utilize the NHP model to evaluate HIV-1 Env-based vaccines.

## Results

### J038 lineage Abs broadly neutralize diverse HIV-1 strains

We previously found that the plasmas from one SHIV_1157ipd3N4_-infected Chinses rhesus macaque (G1015R) neutralized 65% of a panel of 17 tier 2 viruses by week 321 post infection (Gao et al., 2018). The G1015R plasma from week 350 still maintained broad neutralization activity, neutralizing 71% of the same panel of the 17 viruses (Figure 1A). Our previous epitope mapping results showed that many mutations (N160A, T162A and K169E) in the variable loop 2 (V2) in Env rendered the virus resistant to the neutralization by the week 321 plasma, indicating that the V2-specific bnAbs might have been elicited in G1015R (Gao et al., 2020; Gao et al., 2018). To delineate if such bnAbs were elicited in G1015R, we used the peripheral blood mononuclear cells (PBMCs) collected at week 350 to sort 172 single memory B cells that specifically bound to V2 (positive for wild type A244 gp120 but negative for the mutant A244 carrying both N156K and N160K mutations) into individual wells in 96-well plates (Figure 1B).

**Figure 1.**
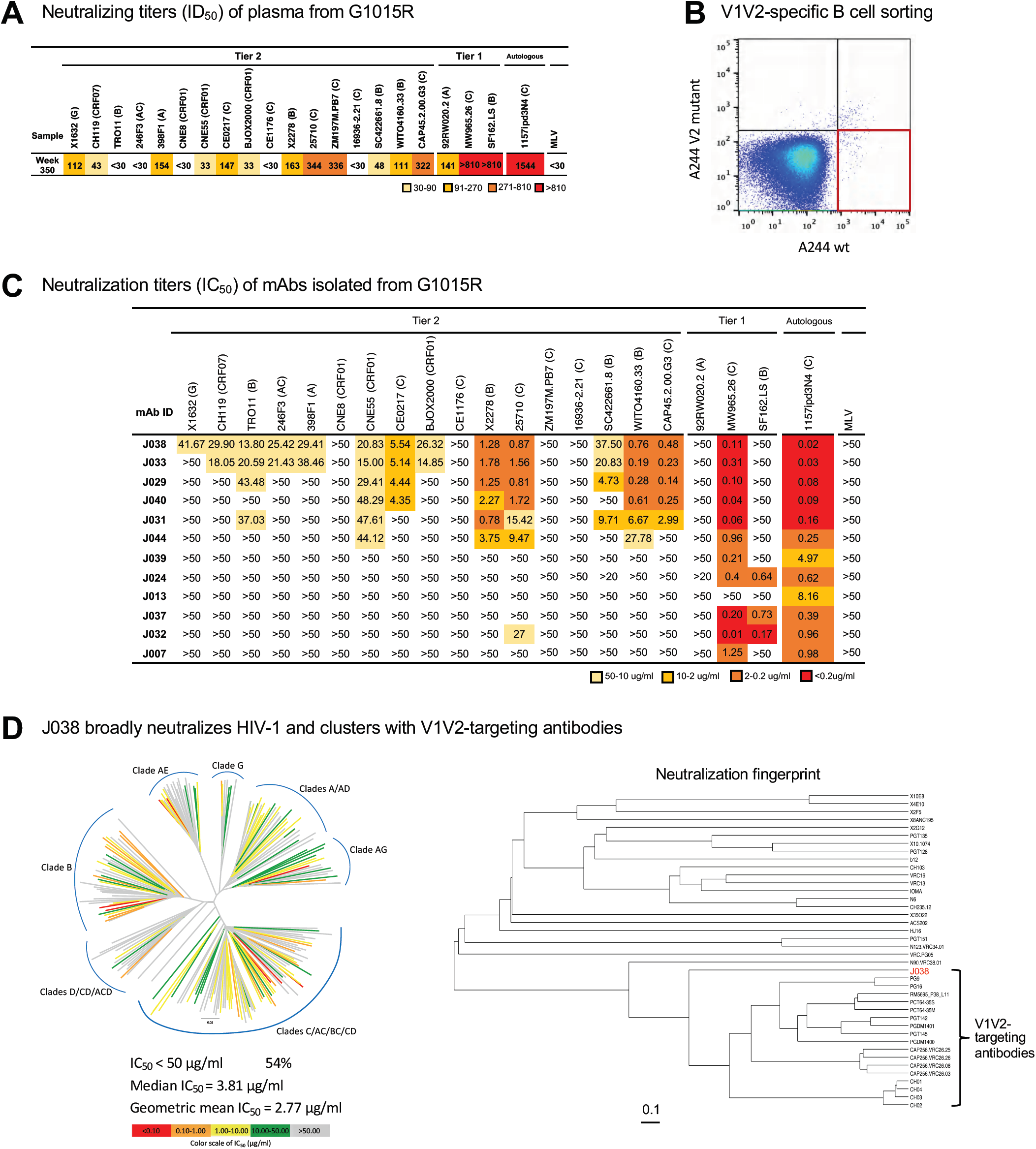
**Isolation of broadly neutralizing antibodies from G1015R** (A) Neutralization breadth of week 350 plasma post infection from G1015R infected with SHIV_1157ipd3N4_. Data are shown as ID_50_ and neutralization potency is color coded. (B) Fluorescence-activated cell sorting (FACS) plots of sorted V1V2-specific single B cells from week 350 peripheral blood mononuclear cells (PBMCs) from G1015R that bind d11_A244_gp120_SavBV421 but not glycan knockout mutant N156_160K_d11_A244_gp120_SavAF647 (red box). (C) Neutralization profiles of 12 mAbs were determined with 17 tier 2 viruses, three tier 1 viruses and the autologous virus. Murine leukemia virus (MLV) was used as a nonspecific control. Neutralization titers are represented as IC_50_ (μg/mL). (D) J038 neutralizes 112 strains of a 208-virus panel (54%). Stains in HIV-1 phylogenetic tree are colored according to their respective J038 potency based on IC_50_. Median and geometric mean titers are calculated only for samples with IC_50_<50 µg/ml. Neutralization profile of J038 shows that it clusters with V1V2-targeting antibodies. **See also Figures S1-S2 and Table S1-S2.**

The paired variable Ig heavy and light chains (VH and VL) were amplified from 44 single cells (Figure S1A). The recombinant protein for each Ab was expressed by co-transfecting the Ig VH and VL genes into the 293T cells. Twelve mAbs showed specific binding to autologous SHIV_1157ipd3N4_ gp120 protein (Figure S1A). Sequence analysis showed that the V(D)J sequences of six Abs (J029, J031, J033, J038, J040 and J044) were from the same gene family (Table S1). Maximum likelihood trees showed that the VH and VL genes from these six Abs evolved similarly, forming two distinct clades; clade A with four sequences and clade B with two sequences (Figure S1B). They all had 18 and 9 amino acids (AAs) in the complementarity-determining region 3 (CDR3) of VH and VL chains, respectively (Table S1). The average of somatic hypermutation (SHM) frequencies were 19.83% and 17% in VH and VL, respectively. All but J044 bound to autologous 1157ipd3N4 gp120 and heterologous BG505 Env trimer at high affinity (Figure S2). The V(D)J sequences in the VH and VL genes had distinct origins in the other six Abs, indicating that they derived from independent germlines (Table S1).

When tested with the same panel of 17 tier 2 viruses, all six members from the same lineage neutralized the viruses at various levels (0.14 to 48.29 µg/ml), with the neutralization breadth ranging from 23.5% to 76.5% (Figure 1C). Among them, two (J033 and J038) of three most evolved antibodies (Figure S1B) showed broadest neutralization activities (70.6% and 76.5%, respectively). The similar neutralization patterns between both mAbs and week 350 plasma suggested that both Abs represented the overall plasma neutralization activity well. In contrast, all other six mAbs did not neutralize any of those 17 viruses, except one case (J032 weakly neutralized 25710). All 12 mAbs potently neutralized autologous tier 2 virus 1157ipd3N4 (Siddappa et al., 2010) and they also neutralized tier 1 virus MW96526, except J013. None of them neutralized tier 1 virus 92RW020.2 and only three of them neutralized tier 1 virus SF162.SL. These results demonstrated that bnAbs representing the overall plasma neutralization activity were induced in G1015R after seven years of infection.

The neutralization potency and breadth of J038 were further assessed with a panel of 208 viruses representing global major circulating HIV-1 strains. Neutralization results from this large panel showed that J038 could broadly neutralize 112 viruses, or 54%, at a maximum of concentration threshold of 50 µg/ml (Figure 1D, Table S2). The median IC_50_ was 3.81 µg/ml. Representative viruses from major subtypes and recombinant forms could be neutralized by J038, demonstrating that the neutralization breadth of J038 is not biased by particular subtypes (Figure 1D). Neutralization fingerprint analysis indicated that the neutralization profile of J038 clusters with those of the V1V2-targeting antibodies, such as PG9, PGT145 and VRC25.26, suggesting its V1V2- specificity.

### Cryo-EM structure reveals J038 Fab binds to the V2-Apex

To determine the structures of J038 or J033 in complex with the HIV-1 Env trimer, we used the neutralization results to select a sensitive strain to make a stabilized trimer (Table S2). The Env from a CRF01_AE strain C1080, which is one of the most sensitive strains to J038 neutralization (IC_50_ of 0.095 µg/ml), was stabilized in the prefusion-closed state using the approach described previously (Rawi et al., 2020). To purify the C1080 Env in native buffer condition, we captured the expressed Env onto a protein A column preloaded with a CD4-binding site antibody 3BNC117, in which an HRV3C cleavage site was engineered into its hinge region for enzymatic release of the antigen-binding fragment (Fab). We then mixed J038 or J033 Fab with the C1080-3BNC117Fab complex released from the protein A column and further purified for electronic microscopy (EM) studies. Initial negative stain EM indicated that one Fab of J033 or J038 bound to each of the Env apex (Figure S3). Cryo-EM analysis with 3-dimensional reconstruction of the J038-C1080-3BNC117 complex at a nominal resolution of 3.4 Å confirmed a single J038 Fab bound to the V2-apex of the C1080 trimer with one 3BNC117 Fab bound at the CD4-binding site of each protomer (hereafter, we refer protomers as P1, P2 and P3, clockwise when viewing down to the Env apex) (Figures 2A-2B, S4 and Table S3). Unlike some V1V2-specific antibodies like VRC26.25 and PGT145, the heavy chain complementarity-determinant region 3 (CDR H3) of J038 did not insert into the trimer interface at the V2 apex. The interface between antibody J038 and the C1080 Env was well resolved with clear density for glycan N160 extending from the surface of Env to interact with the heavy chain CDRs of the J038 Fab. The density of glycan N156 was evident for the portions close to the surface of Env. We also obtained cryo-EM structure for J033, a member of the J038 lineage antibodies. The structure at 4.75 Å resolution showed a very similar binding mode to that of J038 (Figure S4E). We chose the J038 structure that is at higher resolution for further analysis.

**Figure 2.**
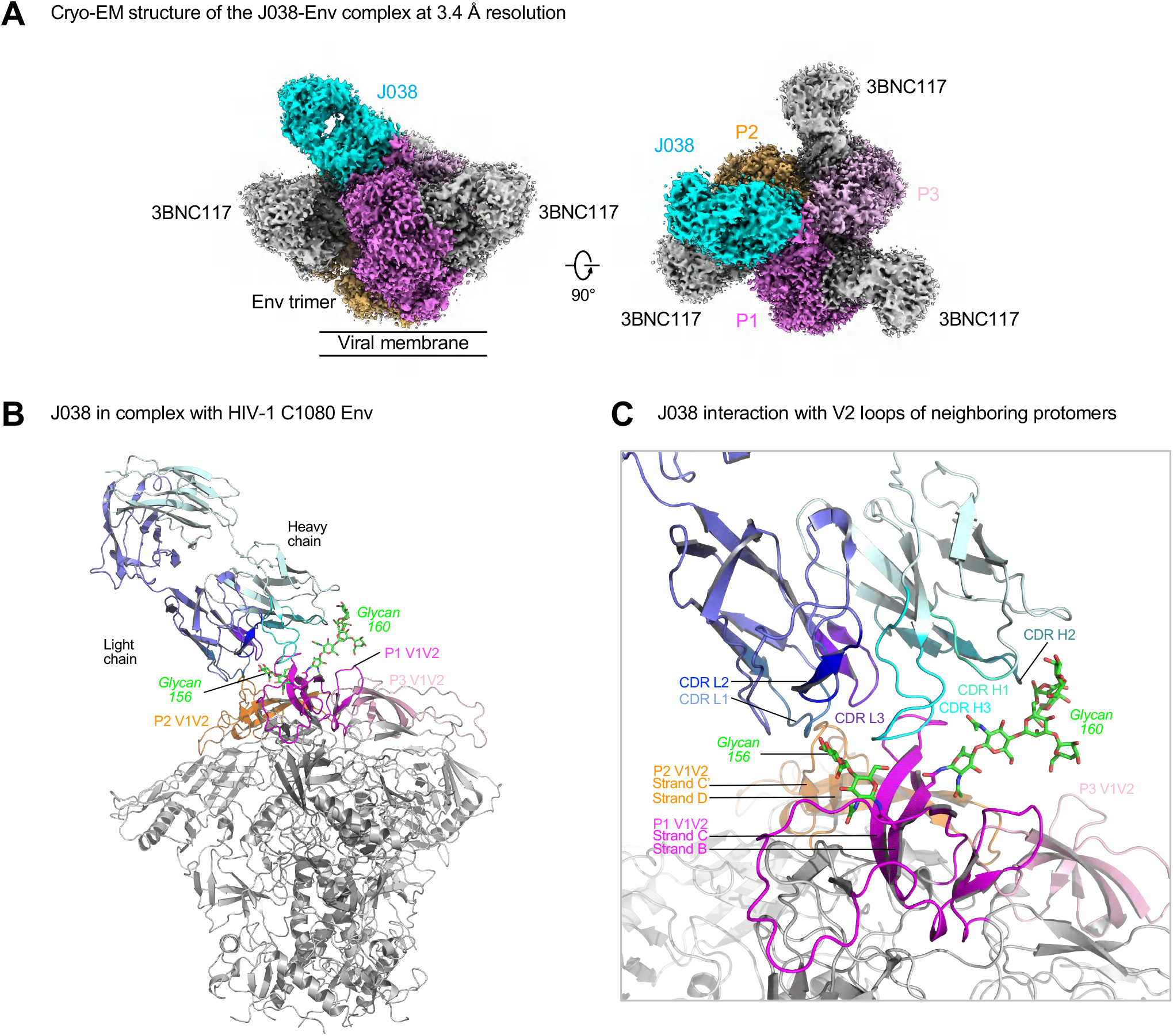
**Antibody J038 targets the V2-apex of the HIV-1 Env.** (A) Overall cryo-EM structure of the J038 Fab -Env complex, with EM reconstruction density shown in gray. The CD4-binding site antibody 3BNC117 was used to aid the resolution. Protomers of the trimer are labeled as P1, P2 and P3. (B) Antibody J038 recognition of the HIV-1 Env. Heavy and light chains of J038 are colored in cyan and blue, respectively. The V1V2 loops of the Env are colored magenta, pink and orange for the three protomers, respectively. Glycans N156 and N160 on P1 interacting with J038 are colored green. The CD4-binding site antibody 3BNC117 was omitted for clarify. (C) J038 interaction with V2-loops of the neighboring protomers P1 and P2. The complementarity determining regions (CDR) of J038 are shown in shades of cyan and blue. The strands of V2 that interacted with J038 are labeled. **See also Figures S5 and Tables S1-S4.**

Overall, the interface between J038 and C1080 Env trimer buried 1298 Å^2^ surface area from the antibody and 1439 Å^2^ from the HIV-1 Env (Table S4A). The epitope of J038 contained both protein and glycan components, of which 89% of the surface was from V2 loop of protomer P1 and 11% of that was from V2 loop of the P2 protomer. Glycans N156 and N160 on strand B of V2 loop on P1 contributed 45% of the total epitope surface (Figure 2C and Table S4A). On the antibody side, both heavy chain and light chain were involved in binding of the V2 apex. While heavy chain, which contributed 75% of the paratope, interacted with protomer P1 residues and glycans, light chain contacted both protomers P1 and P2 (Figure 2C and Table S4A)

### J038 interacts with the V1V2 B-strands on two protomers of the Env

All CDRs of J038 contributed to binding of C1080 Env (Figure 2C). Unlike the CDR H3 in antibodies such as VRC26.25 and PGT145 which insert into the V2-apex hole on the Env trimer, the 18-residue long CDR H3 of J038 formed extensive anti-parallel β-strand (main chain) hydrogen bonding interactions with the V2 C-strand of protomer P1 (Figure 3A and Table S4B). Residues at the tip (D99-D100-F100A-Y100C; antibody residues follow Kabat numbering system) of CDR H3 formed hydrogen bonds with side chains of K169, K171 and Y173 in the same C strand. In addition, heavy chain residues Y33 in CDR H1 and R54 in CDR H2 formed hydrogen bonds with D167 and R166 located in the loop between strands B and C of protomer (Figure 3A). On the light chain side, the CDR L3 tip residues Y91-R92-R93 formed four hydrogen bonds with E164 and K168 on the same loop in protomer P1 (Figure 3A, middle). We also observed hydrogen bonding interactions between J038 light chain residues D1, CDR L1 R27 and CDR L3 R93 and protomer P2 residue N186 located in the loop between V1V2 strands C’ and D (Figure 3A, right). Although J038 is a V2-Apex bnAb, its EDDY motif mutated into a EDDF motif that interacted with L171 and Y173 not A166 and L169 as other V2-Apex bnAbs (Gorman et al., 2020; Gorman et al., 2016; Lee et al., 2017; McLellan et al., 2011), suggesting that J038 binding to the V2-Apex distinctively.

**Figure 3.**
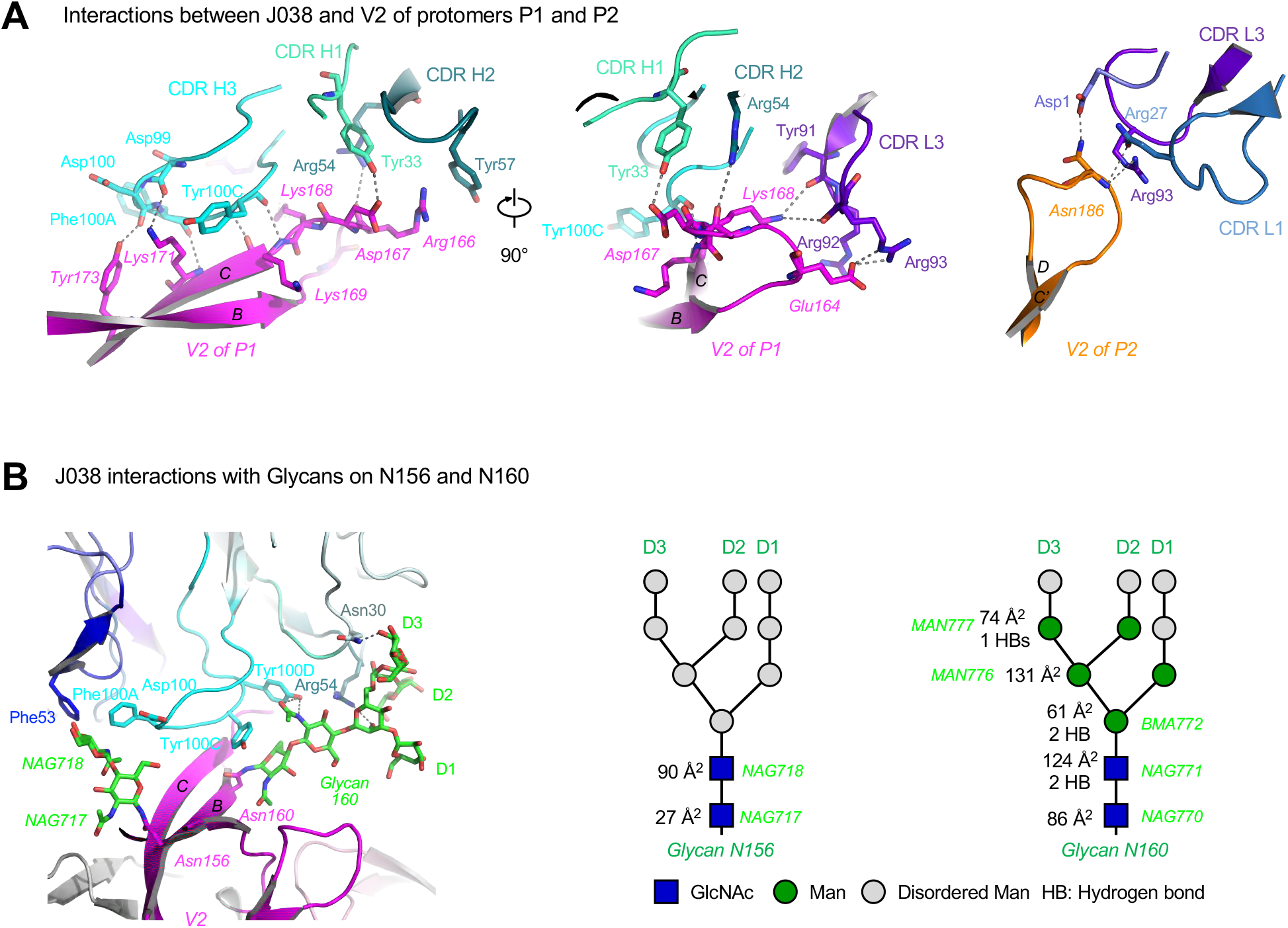
**J038 interaction with V2-apex of HIV-1 Env.** (A) J038 interaction of V2 of protomers P1 (left and middle) and P2 (right). Key interacting residues are shown in sticks and numbered with hydrogen bonds shown in dashed lines. (B) Interactions between J038 and glycans on Asn156 and Asn160. Key residues that interact with glycans are shown in sticks and numbered with hydrogen bonds shown in dashed lines. A schematic for each glycan are shown next to the cryo-EM structure with GlcNAc as blue squares and mannose residues as green circles. Buried surface areas (Å^2^) and hydrogen bonds with J038 for each glycan moieties are labeled. **See also Table S3-4.**

### The binding of J038 to Env is heavily dependent on glycans

Similar to other V2-Apex bnAbs, J038 interacted with V2-Apex through extensive binding to glycans (Figure 3B). The N160 glycan accounted for the majority (80%) of the total glycan contact (Figure 3B and Table S4A). Through the two base GlcNAc, BMA and the Mannose moieties on the D3 branch, glycan N160 interacted with J038 heavy chain by forming 5 hydrogen bonds with N30, R54 and Y100D in all three CDRs (Figure 3B and Table S4B). These interactions accounted for over one-third (37%) of the total epitope area. The N156 glycan is responsible for 8% of the total antibody-Env contact area. Of the two ordered N156 glycan GlcNAc moieties, NAG717 only interacts with D100 in CDR H3 while NAG718 binds to F53 in CDR L2 as well as D100 and F100A in CDR H3. Overall, the glycan contributed nearly half (45.5%) of the total J038-binding surface, indicating that the glycans plays a very important role in binding of J038 to V2-Apex.

Viral sequence alignments showed that C1080 and SHIV_1157ipd3N4_ share the identical sequences on stands B and C of the V2 loop (Figure 4A). This explains why C1080 is potently neutralized by J038 (Table S2). The N160 glycan in SF162 was lost due to the N160K mutation and this rendered SF162 full resistant to the J038 lineage bnAbs. Even though the loop between stands C’ and D is slightly divergent between strains in length and residue composition, the four epitope residues (E185, N186, K187 and N189) in this region on protomer P2, which contacted CDR L1 and L3 regions of J038 and provided 11% of the epitope surface, were relatively conserved (Figure 4A, Table S4B). Overall, the structural information revealed in the J038-C1080 complex provided structural basis for its breadth of neutralization.

**Figure 4.**
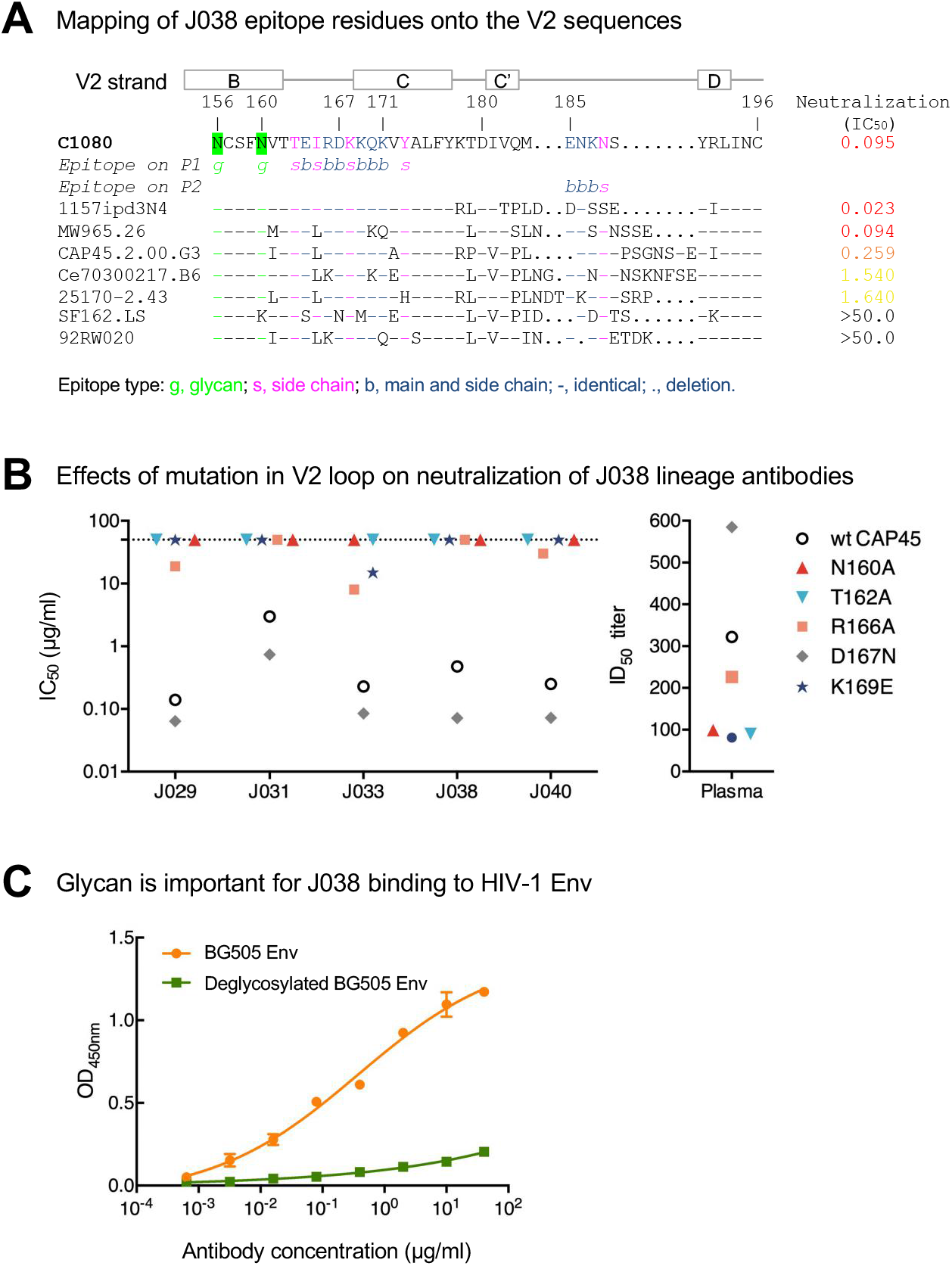
**Key J038 residues interacting with V2-apex of HIV-1 Env.** (A) Alignment of V2 sequences from HIV-1 strains, shown relative to their neutralization sensitivities. J038 contacts are marked as g for glycan contacts, s for side chain contacts and b for both side chain and main chain contacts. (B) Effects of mutations in V2 on neutralization potency of the J038 lineage antibodies. Mutations removing the Asn160 glycosylation site on V2 make the virus resistant to J038 lineage antibodies. (C) Deglycosylation reduces binding of HIV-1 BG505 Env to J038. **See also Figure S5 and Table S4.**

To confirm the requirements of key residues and glycans on neutralization, we performed mutagenesis studies with CAP45, a strain potently neutralized by 5 of 6 members of the J038 lineage antibodies (Figure 1C). When the glycosylation site at position 160 was eliminated by either N160A or T162A mutation, CAP45 became fully resistant to neutralization by all five J038 lineage members (Figure 4B). Two other mutations (R166A and K169E) rendered CAP45 more resistant potentially by removing key interactions, such as the stacking effect with Y57 in CDR H2 (Figure 3A and 4B), but the D167N mutation made CAP45 slightly more sensitive. Similar results were observed with the week 350 plasma from G1015R (Figure 4B). Deletion of the N160 glycosylation site in two other viruses (25710 and Ce0217) also rendered the viruses full resistant (Fig. S5A). When deglycosylated BG505 trimer was used, the binding of the J038 lineage members were dramatically reduced (Figure 4C and S5B). These results showed that N160 glycan and some key contact sites are required for efficient binding of the J038 lineage antibodies to V2-Apex.

### J038 binding induces V2 into a unique “Up” position

Most V2-Apex bnAbs, such as PG9, VRC26 and PGT145, use their long CDR H3 to penetrate into the apex hole formed at the V2-Apex and interact with the entrance regions of strand C through extensive hydrogen bonding (Gorman et al., 2020; Gorman et al., 2016; Lee et al., 2017; McLellan et al., 2011). In contrast, J038 and J033, which had short CDR H3, bound to the V2-apex without penetrating into the apex hole in a similar manner to VRC38 (Figure 5A). Interestingly, when J038 bound to the Env trimer, it induced the V2 strands B and C in protomer P1 into an “up” position (Figure 5B). This unique antibody-induced V2 “up” position has not been observed in any other V2-Apex bnAb/Env complex (Figure 5C). When aligned over the whole Env trimer, the J038-bound V2 strands B and C (residues 154-177) in protomer P1 had an average C*α*-RMSD of 7.2 between corresponding regions in other antibody-bound Env (Figure 5C). In the J038-bound mode, the tip (C*α* of residue 165) of the loop between B and C in protomer P1 moved 11.6 Å to 18.7 Å away from the apex compared to positions in other antibody bound forms (Figure 5D). The strands B and C in the other two gp120 protomers had very similar conformation between the J038- and other antibody-bound Env, for example, that in P2 between J038- and PGT145-bound only had a RMSD of 0.7 Å. In summary, the J038-induced “up” position of V2 loop represent the first-time observation of such a unique and asymmetric conformation in HIV-1 Env. This unique “up” position of V2-Apex allows easy and tight interaction with CDR3 H3 of J038 (Figure 2C and 5A).

**Figure 5.**
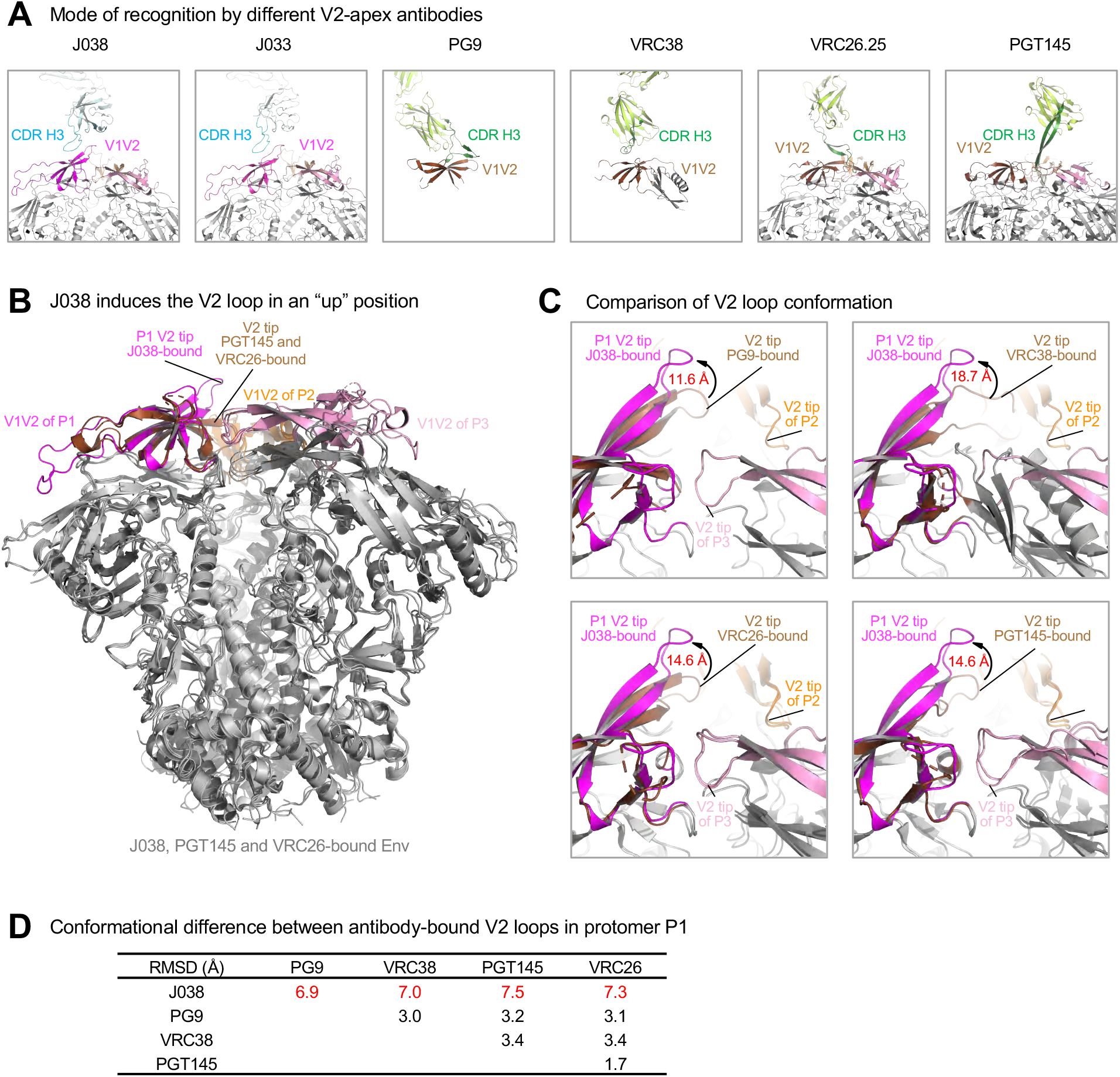
**J038 uses a different mode of gp120 recognition.** (A) Comparison of heavy chain CDR H3 interactions with the V2-apex of HIV-1 Env. CDR H3s of antibodies (J008, J033, PG9, VRC38, VRC26 and PGT145) are shown in cyan or dark green. The V1V2 loops are colored as in Figure 1 with that of protomer 1 in the PG9-, VRC38-, VRC26- and PGT145-bound shown in brown for comparison. (B) Binding of J038 induces the strands B and C in P1 V2-loop in an “up” conformation. The structures of J038, VRC26 and PGT145 in complex with HIV-1 Env are superposed and the V1V2 loops are colored the same as in (A). (C) Comparison of conformation of the P1 V2-loops in J038, PG9-, VRC38-, VRC26- and PGT145-bound structures, distance shown are between C*α* atoms of Glu165 at the tip of loop between strands B and C. (D) C*α*-rmsd between antibody-bound V2 loop strands B and C (residues 154-177) in P1.

### J038 maturation through gradually increased binding areas to glycans

To understand how the J038 lineage Abs evolved to more mature overtime, we deduced the unmutated common ancestor (UCA) for both VH and VL sequences and inferred the intermediate antibodies (IAs) (Figures 6A and S1B). There are two clades of the sequences; clade A included all four members (J029, J033, J038 and J040) which are broadly neutralizing, and clade B contained two members (J031 and J044) that neutralized relatively fewer heterologous tier 2 viruses (Figure 1C). To investigate how the gradually accumulated mutations affect neutralization breadth and potency of the J038 lineage Abs, we synthesized and expressed the recombinant UCA and all five IA antibodies. The UCA bound to the heterologous BG505 Env very weakly but gradually gained high affinity as it became more matured (Figure S6). However, like the CH103 lineage Abs (Liao et al., 2013), the UCA bound the autologous 1157ipd3N4 Env well and the binding affinity of matured J038 was only about 5-fold better than the UCA (Figure S6).

**Figure 6.**
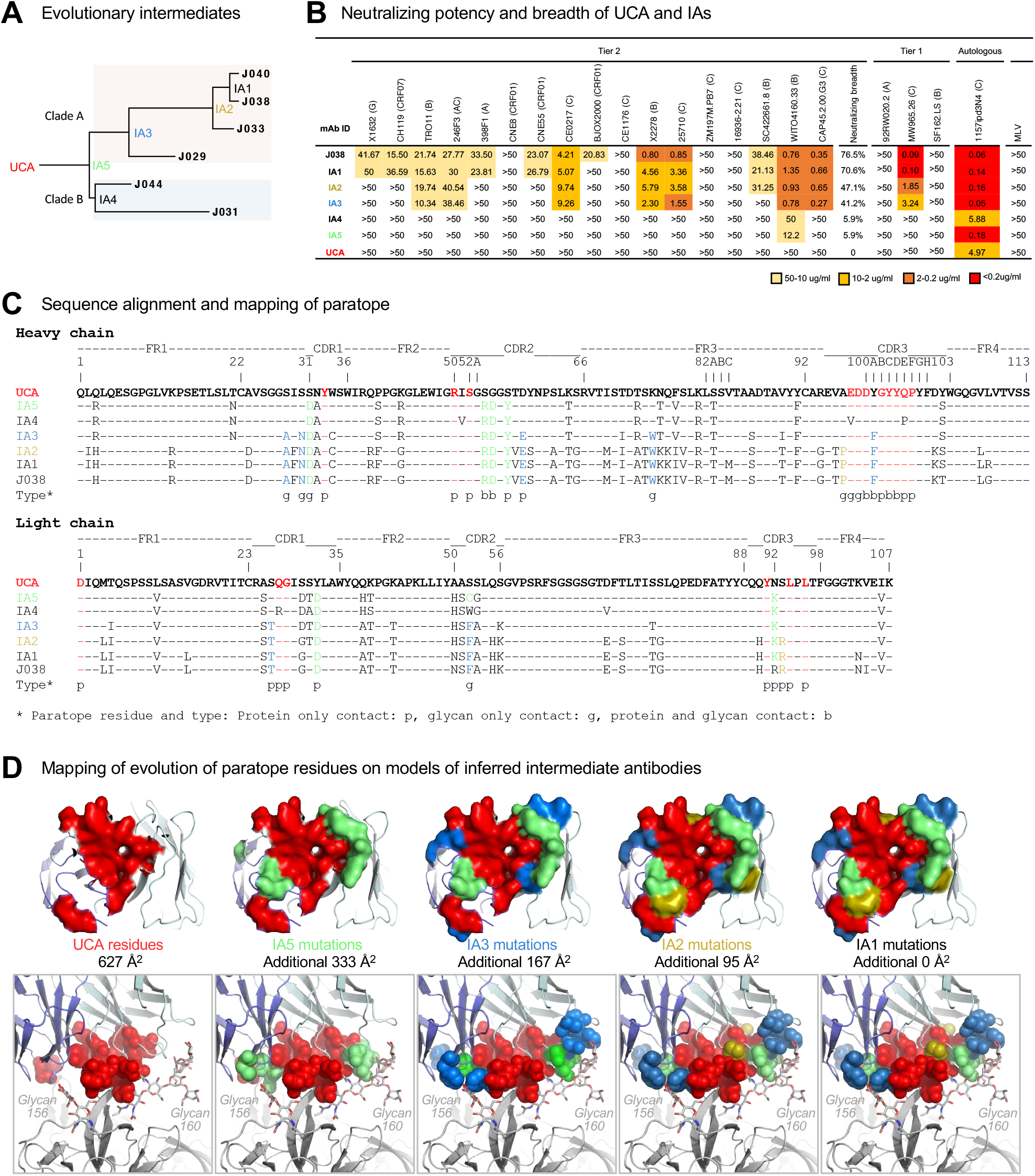
**Development of neutralization breadth during maturation of the J033 lineage Abs.** (A) The maximum likelihood phylogenetic tree of the J038 lineage sequences. UCA and IAs at evolutionary nodes were computationally inferred. Two different clades are shaded with different colors. (B) Neutralization profiles of UCA and IAs were determined with 17 tier 2 viruses, three tier 1 viruses and the autologous virus. Murine leukemia virus (MLV) was used as a nonspecific control. Neutralization titers are represented as IC_50_ (μg/mL). (C) Sequence alignment of the UCA, intermediates and mature J038 Ab sequences. Heavy chain shown in the upper set and light chain shown in the lower set. Env contacts are shown as g (glycan contacts), s (side chain contacts) and b (both side chain and main chain contacts). (D) Mapping of the development of J038 paratope residues on models of inferred intermediate antibodies. Paratope residues directly from UCA were colored red, those from IA5, IA3 and IA2 were colored green, light blue and olive, respectively. Paratope surface areas estimated from UCA and increase from intermediate antibodies are indicated. Top panels: on antibody. Bottom panels: interface of antibody and V2-apex. **See also Figures S7 and Table S5.**

When tested against the same 22-virus panel, the UCA did not neutralize any of the tier 1 and 2 viruses, but it neutralized autologous tier 2 1157ipd3N4 pseudovirus well at 4.97 µg/ml (Figure 6B). The least evolved intermediates IA5 and IA4 were similar and only weakly neutralized one tier 2 virus WTO4160.33 and autologous virus 1157ipd3N4. However, IA5 more potently neutralized the autologous virus than UCA while IA4 that led to weakly neutralizing clade B Abs maintained the same low neutralization level as UCA. IA3 and IA2 became more potent by neutralizing 7-8 strains of tier 2 viruses (39%-44%) and one of three tier 1 viruses. IA1, which neutralized 12 tier 2 viruses, gained even more neutralization breadth (70.6%) similar to that of J038, missing only one virus (BJOX2000) (Figure 6B and 1C). Compared to IA5 and IA4, both IA3 and IA2 became 17-fold more potent to neutralize tier 1 virus MW965.26. However, J038 is 2.3 folds more potent to neutralize autologous virus than IA1. Interestingly, once the lineage Abs gain the ability to neutralize a heterologous tier 2 virus, the level of the neutralization against the virus remain the same during the Ab maturation process. Taken together, these results demonstrate that the neutralization breadth of the J038 lineage Abs gradually increase as they mature over time.

To understand the relationship between J038 maturation and the development of breadth, we compared the sequences of UCA, intermediate and mature antibodies to identify the somatic hypermutations along with the paratope residues of J038 (Figure 6C). We also performed template-based homology modeling of the UCA and intermediate antibodies using the J038-C1080 structure (Figure 6D). Mapping of the paratope indicated that 11 and 6 of the paratope residue were directly encoded by the VH and VL genes of UCA, respectively, which provided 627 Å^2^ binding area (Figures 6C-6D and Table S5A). In IA5, four and three residue substitutions in VH and VL, respectively, evolved at the contact sites and added an additional binding area of 333 Å^2^. IA3 gained another 4 and 2 residue substitutions in VH and VL, respectively, at the contact sites, and the binding area was increased by another 167 Å^2^. When there was only one residue substitution was found in VH or VL region of IA2 and both together further increased binding area by 95 Å^2^. The additional mutations in IA1 and mature Abs did not directly alter the binding area (Figures 6C-6D and Table S5B).

We observed that paratope residues from UCA primarily provided core interactions to stand C of protomer P1 (Figure 6D). The early increased binding area by IA5 further strengthened the contact with protein components of V2 strand C. It is interesting to noticed that the later increased binding area by IA3 and IA2 were at the peripheral of the paratope and were mainly (6 of 10 residue substitutions) for binding of the N156 and N160 glycans. These results demonstrate that the J038 lineage Abs matured by strengthening interactions to the V2 residues and then increasing their binding areas to glycans on Env to broaden the neutralization breadth (Figure 6D).

We also performed Biolayer interferometry (BLI) assay to determine the binding kinetics of inferred antibodies (Figure S6). The affinity of UCA and IAs to autologous gp120 was consistent with their neutralizing potency against the autologous pseudovirus. The BLI assay also showed that IA4, the progenitor for clade B Abs in the phylogenetic tree, bound to HIV-1 Envs much weaker than IA5, similar as UCA. J031 and J044 were evolved from IA4 and both had a narrower neutralization breadth (Figure 1D). These results indicate that the mutations accumulated in IA4 had a negative influence on antibody evolution towards broad neutralization.

### Limited mutations at the Ab-Ag interface in the *env* gene

A small region (positions 156-188) including two glycosylation sites (N156 and N160) in strands B and C of V2 was responsible for the contact with J038 (Figure 3). Examination of viral sequences at this region from the longitudinal plasma samples in G1015R showed no mutations at week 27. No plasma samples from the next 187 weeks were available for sequence analysis. By week 214, all 23 viral sequences had the I165L and K171R mutations, while the K168R and V172A mutations accounted for 69% and 92% of the virus population, respectively (Figure 7A). All three mutations (I165L, K171R and V172A) are fixed in the viral population by week 350 while the K168R mutation became undetectable. Four residues in the V2 strands C’ and D from protomer P2 in the Env trimer were found to contact J038 (Figure 4A). However, no mutations in this region were detected by week 214 (Figure 7A). A few different mutation combinations were found in some sequences by week 350. Interestingly, no mutations that can eliminate N156 and/or N160 glycosylation sites were found throughout the 7-year follow-up period.

**Figure 7.**
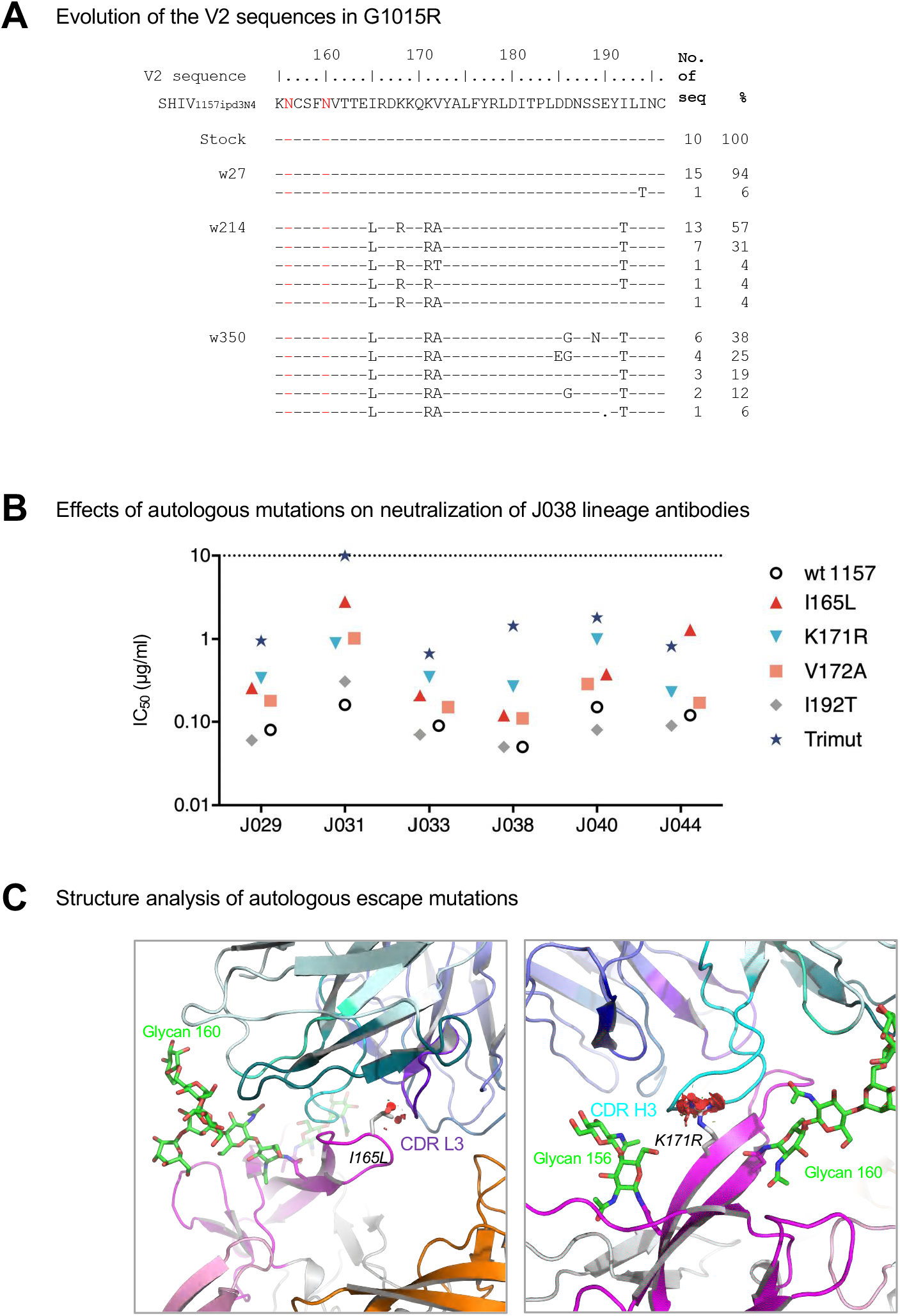
**Limited mutations accumulated at the J038-Env contact site during infection.** (A) Identification of neutralization escape mutations in the *env* gene from longitudinal samples in G1015R. Amino acid sequences of V2 from the viral stock and weeks 27, 214 and 350 post infection were compared to the SHIV_1157ipd3N4_ reference sequence. The amino acid positions are based on the HIV-1 HXB2 sequence. (B) Effects of the mutations in V2 on neutralization potency of the J038 lineage antibodies. I165L, K171R and V172A render the virus resistant to J038 lineage antibodies at various levels. (C) Structural basis for neutralization resistance by the I165L and K171R mutations. Leu165 and Arg171 are shown in stick representation. Potential clashes with CDR l3 and CDR H3 caused by these mutations are indicated as red discs.

To understand how these mutations affected viral susceptibility to neutralization by the J038 lineage antibodies, we introduced these three mutations (I165L, K171R and V172A) individually or all together into the autologous 1157ipd3N4 *env* gene. Since both the I165L and K171R were the Env-J038 contact sites, the I165L and K171R mutants were 7 and 5 folds more resistant, respectively, to the J038 lineage Abs (Figure 7B). The V172A mutation was not a contact site and the V172A mutant was only slightly more resistant to neutralization (3 folds) than the wild type 1157ipd3N4. However, when the I165L, K171R and V172A mutations (Trimut) were tested together, it was about 20-fold more resistant. We also tested the I192T mutation which is not at the Env-J038 contact site but fixed in viral population by week 214 and 350. The I192T mutant had similar neutralization susceptibility as the wild type 1157ipd3N4.

Structure modelling of the I165L and K171R mutations at the contact sites showed that both mutations might potentially cause clashes with CDR L3 and CDR H3, respectively (Figure 7C). This explained why both mutations alone modestly increased resistant to neutralization by the J038 lineage antibodies. Although V172A is not involved in antibody binding, but all I165L, K171R and V172A mutations together could have a collective unfavorable interaction with J038 (Figure 7C). These results demonstrate that over seven years of infection, a few mutations accumulated at the J038 contact sites in the *env* gene and only modestly impacted viral susceptibility to neutralization by the J038 lineage Abs in G1015R.

## Discussion

We sorted single memory B cells from SHIV-infected Chinese rhesus macaques using HIV-1 Env V2-Apex differential baits and obtained bnAbs that can broadly neutralize 54% of the 208 HIV-1 strains. This is the first case to show that bnAbs can be successfully obtained from monkeys that developed broad neutralization activity in sera after seven years of infection. This demonstrates that the bnAbs can be elicited in SHIV infected macaques after a long period of infection time as determined by longitudinal plasma samples and reconstructed Abs evolved from UCA to mature bnAbs. This lineage of bnAbs targets V2-Apex in the HIV-1 Env trimer. However, sequence, structure and phenotype analyses of the J038 lineage bnAbs show a number of unique features that can have important implications in future HIV vaccine design.

The six J038 linage bnAbs are the only predominant gene family (14%) among 44 sorted B cells. None of other 38 Abs share the same VH and VL sequence from the same gene family. The J038 lineage bnAbs were derived from VH4 gene family, which is dominant usage in rhesus macaque B cell repertories (Dai et al., 2015; Sundling et al., 2012a). A number of nAbs obtained from macaques are found to belong to the VH4 gene family (Dai et al., 2015; Han et al., 2019; Kong et al., 2019; Sundling et al., 2012a; Wang et al., 2017). These results indicate that the VH4 gene family may be preferred for bnAbs, regardless any binding specificities, in SHIV infected or vaccinated macaques. Similar VH germline orthologous gene is also found in humans (Figure S7), indicating that such Abs are likely to be induced in humans too.

Most of previous V2-apex targeting bnAbs obtained from HIV-1 infected humans have a long anionic CDR H3 and high SHM except VRC38 (Andrabi et al., 2015; Cale et al., 2017; Gorman et al., 2016; Moore et al., 2017). However, the CDR H3 of the J038 lineage bnAbs, like VRC38, are short, with only 18 amino acids. Like other V2-apex bnAbs (Gorman et al., 2020; Gorman et al., 2016; Lee et al., 2017; McLellan et al., 2011), only one J038 Fab binds to the Env trimer. However, binding of J038 induces a unique “up” conformation of V2 in the main binding protomer, which provides ∼90% of the epitope surface, while the other two V2 loops in the same Env trimer remain unchanged. J038 binding to Env is heavily dependent on two glycans (N156 and N160), accounting for 45% of the total contact area. The deletion of either of these glycosylation sites in heterologous *env* genes results in much reduced binding and renders the viruses resistant to neutralization. The “up” position of V2 induced by J038 represents the first observation of such conformation in HIV-1 Env not identified in other V2-apex bnAb-bound Envs (Cale et al., 2017; Gorman et al., 2020; Lee et al., 2017; Walker et al., 2009). This unique binding pattern of J038 may facilitate tighter interaction with J038 and thereby enables binding to monomeric gp120.

Compared to the other bnAbs with V2 or other specificities (Klein et al., 2013; Kwong et al., 2013; Sok and Burton, 2018), the J038 linage bnAbs does not have strong selection pressure on the viral populations during 7 years of infection. Only three mutations (I165L, K171R and V172A) at the Ab-Env contact sites were strongly selected by week 350 post infection. Among them, the I165L and K171R mutations interact with J038 and either mutation rendered the virus slightly more resistant to neutralization by J038. No mutations that affect either N156 or N160 glycosylation sites were detected. This suggests that the maturation of neutralization breadth in G1015R may not require extensive mutations at the contact sites.

To increase the neutralization breath, the CH235 lineage bnAbs decrease the binding area size to focus on only the highly conserved CD4 binding site core on Env (Bonsignori et al., 2016) while the CH103 lineage bnAbs change the angle of VL to more likely accommodate the size changes of the V5 loop during their maturation process (Fera et al., 2014). The J038 lineage bnAbs gain its neutralization breadth through a different mechanism. On the core binding area contacted by UCA-encoded residues, the mutations in more evolved intermediary IA5 increased 54% of the binding area by mutations in both VH and VL chains. More mutations accumulated in IA3 and IA2 further increased the binding by another 42% binding area. The increased binding area between Ab and Env during the maturation process of the J038 lineage bnAbs allow them to bind Env at higher affinity. Importantly, the majority increased binding from UCA is through binding to both N156 and N160 glycans (mainly N160 glycan).

Our previous studies showed that the UCA of bnAbs do not neutralize T/F viruses or heterologous viruses that initiated the bnAb lineage (Bonsignori et al., 2017; Gao et al., 2014; Liao et al., 2013). However, the UCA of DH727 obtained from a vaccinated macaque could neutralize a heterologous CNE8 (Han et al., 2019), and the UCA of CAP256-VRC26 could neutralize CAP256-SU weakly (Doria-Rose et al., 2014; Gorman et al., 2016). Here we found that the UCA of J038 lineage antibodies moderately neutralize autologous 1157ipd3N4 virus, but not any other tier 1 and tier 2 viruses. This demonstrate that the germline of some bnAbs is fully capable to neutralize autologous T/F viruses. Thus, bnAbs derived from such germlines may be easier to induce by vaccines than those bnAbs derived from germlines that cannot neutralize the autologous T/F viruses that trigger the lineages.

When the J038 lineage bnAbs continue to evolve, they become more potent to neutralize the autologous virus and gain the ability to neutralize more heterologous tier 2 viruses. Since N156 and N160 glycans are conserved among global HIV-1 strains and both contribute nearly 50% of the Ab binding area, the increasing binding area to both glycans may lead to broad neutralization of diverse HIV-1 strains. Importantly, the invariant of both glycans during 7 years of infection allows the J038 lineage bnAbs to continuously gain additional binding area to them during their maturation process.

In summary, mature J038 is broadly neutralizing (54% of HIV-1 stains), the J038 lineage is the predominant V2-binding Abs in G1015R, the UCA of the J038 lineage bnAbs can neutralize the autologous virus, it binds a unique “up” V2 position, and its maturation heavily depends on glycans. Moreover, J038 has a relatively short HCDR3 and relatively low SHM rate (∼20%). Finally, its maturation does not drive extensive mutations in the Ab-Env contacts sites. All these together indicate that such bnAbs may be easier to induce. This has important implications in HIV vaccine design to induce J038 like bnAbs in humans.

### Limitations of the Study

Broadly neutralizing antibodies with only V2 specificity were characterized using the HIV-1 Env V2-Apex differential baits in this study. This may not fully represent all the bnAb activities, since nAbs with CD4bs and V3 specificities were also identified in the same macaque. HIV-1 Env specific memory B cells were sorted and characterized from only one macaque. Studying additional macaques infected with SHIV_1157ipd3N4_ can further strengthen our findings. Since this study was carried out with the previously archived blood samples, many samples were exhausted for our previous studies. We did not have plasma samples available from week 27 through week 214 to study if viral mutations during this long infection period have any impacts on viral escape from the J038 lineage bnAbs. Since J038 sequence is very similar to its UCA and IA sequences and directly evolved from them, we used the template-based homology modeling to determine the structure basis for the J038 maturation process. Resolving cryo-EM structures between the HIV-1 Env and the UCA and IAs of the J038 lineage bnAbs could reveal structure details in higher resolution.

## Acknowledgments

We thank M. Anthony Moody, Kevin O. Saunders and Barton F. Haynes for providing V2 differential baits and rhesus macaque antibody expression vectors. We also thank members of the Structural Biology Section, Vaccine Research Center, for discussions and comments on the manuscript. This project was supported by the National Natural Science Foundation of China (Grant No. 31970888 and No. 82002138), Key Projects in the National Science & Technology Pillar Program in the Thirteenth Five-year Plan Period (Grant No. 2018ZX10731101-001-010 and 2018ZX10731101-002-003), Program for JLU Science and Technology Innovative Research Team (grant no. 2017TD-05). This research was, in part, supported by the Intramural Research Program of the Vaccine Research Center, National Institute of Allergy and Infectious Diseases (NIAID) and the National Cancer Institute’s National Cryo-EM Facility at the Frederick National Laboratory for Cancer Research under contract HSSN261200800001E.

## Author contributions

Conceptualization, N.G., C.Q., T.Z., X.Y. and F.G.; Data acquisition, N.G., Y.G., L.M., C.W., W.W., X.L., T.G., M.L., N.D.; Data analysis, N.G., C.W., M.L., N.D., B.Y., H.L., C.Q., P.D.K., J.R.M., T.Z., X.Y., F.G.; Bioinformatic analysis, K.W., T.B.K., J.L.; Monkey experiment, W.W., C.Q., H.N.; Neutralization fingerprint analysis A.J., Y.T.; Design and generation of HIV-1 Env proteins, J.G., A.S.O., R.R., P.K.; Cryo-EM analysis, N.D., W.W., C.S., K.H.; Structure analysis, T.Z., P.D.K.; Writing – Original Draft, N.G., Y.G.; Writing – Review & Editing, N.G., X.Y., T.Z., F.G.; Supervision, F.G.; Funding acquisition, C.W., ., F.G

## Supplemental Materials

### KEY RESOURCES TABLE

### RESOURCE AVAILABILITY

#### Lead Contact

Further information and requests for resources and reagents should be directed to and will be fulfilled by Feng Gao.

#### Materials Availability

Expression plasmids generated in this study for expressing antibodies and Env mutants will be shared upon request.

### EXPERIMENTAL MODEL AND SUBJECT DETAILS

#### Cell lines

HEK 293T Cells (Cat# CRL-3216) were from ATCC. Expi293F™ Cells (Cat# A39240) were from Thermo Fisher Scientific. TZM-bl Cells were from NIH AIDS reagent program. 293-6E cells were from NTCC.

#### Macaque PBMC samples

Peripheral blood mononuclear cells (PBMCs) were collected from the Chinese rhesus macaques G1015R infected with SHIV_1157ipd3N4_ via the intrarectal route (Gao et al., 2018). Monoclonal antibodies were isolated from the PBMCs collected at week 350 when broad neutralization activity was detected and the viral load was maintained at a high level.

### METHOD DETAILS

#### Epitope-specific single B cell sorting

Single B cell sorting was performed as described previously (Wiehe et al., 2014; Zhang et al., 2016). Frozen PBMCs (∼1 × 10^7^ cells) were thawed in RPMI 1640 with 10% FBS and washed with PBS with 5% FBS. Multicolor staining was performed using a panel of fluorophore-labeled antibodies (CD3-PE-CF594, CD16-PE-Cy5, CD14-PE-Cy7, CD20-BB515, IgD-PE, CD27-APC-Cy7, AmCyan-DEAD, wild type gp120-BV421, mutant gp120-AF647). All staining was followed by washing and resuspension with PBS with 5% FBS. Single epitope-specific cells were sorted into individual wells in 96-well PCR plates. Each well contains 20 µL lysis buffer consisting of 0.5 U RNase OUT (Invitrogen, Grand Island, NY), 0.0625 µL Igepal (Sigma, St. Louis, MO), 5 µL 5x SuperScript III First-Strand Buffer and 1.25 µL DTT provided with the Superscript III Reverse Transcriptase kit (Invitrogen, Grand Island, NY). The PCR plates containing lysed single cells were stored at -80℃. Sorting data was analyzed with FlowJo software (FlowJo, Treestar, Ashland, OR).

#### Antibody sequence analysis

Single cell RT-PCR for amplification of IgG heavy and light chain genes was performed as described previously (Sundling et al., 2012b; Wiehe et al., 2014). The cDNA of Ab mRNA was synthesized with 150 ng Random Hexamers (Qiagen, Germantown, MD), 200 U Superscript III and 1 mM dNTPs by incubating at 42°C for 10 minutes, 25°C for 10 minutes, 50°C for 60 minutes and 94°C for 6 minutes. The heavy and light chains variable regions were amplified separately from the cDNA templates with two rounds of PCR using panels of specific primers to distinguish different Ab gene families. In the first round PCR, 3.5 μL of cDNA, 5 µL 10× High Fidelity PCR Buffer, 1 µL 10 mM dNTPs, 2 µL 50mM MgSO4, 1 µL 5’-primer mix and 1 µL 25 μM of 3’-primer, 0.25 µL High Fidelity Taq (Invitrogen, Grand Island, NY) were added into a 50 µL reaction. The PCR thermocycling conditions were as follows: one cycle at 94°C for 2 minutes; 35 cycles of a denaturing step at 94°C for 15 seconds, an annealing step at 64°C (Igκ and Igλ) or 62°C (IgH) for 30 sec, and an extension step at 68°C for 1 minute; and one cycle of an additional extension step at 68°C for 5 minutes. The second round PCR was performed with 1 μL of the first round PCR product, 5 μL 10× PCR Buffer, 1 µL 10 mM dNTPs, 1 μL Jumpstart Taq (Sigma, St. Louis, MO) and 1 µL 25 μM of IgH, Igκ, or Igλ inner forward and reverse variable region primers. The second round PCR thermocycling conditions were as follows: one cycle at 94°C for 5 minutes; 35 cycles of a denaturing step at 94°C for 30 seconds, an annealing step at 64°C (Igκ and Igλ) or 62°C (IgH) for 30 sec, and an extension step at 72°C for 1 minute; and one cycle of an additional extension step at 72°C for 10 minutes. PCR products with correct sizes were directly sequenced. Alignments of sequences to germline V, (D), and J genes were performed using IMGT/V-QUEST (www.imgt.org).

#### Expression of recombinant antibodies

Linear Ab expression cassettes were generated with paired heavy and light chain PCR products from the same single cells by adding a CMV promoter to each Ab chain to express recombinant Abs (Liao et al., 2009). When the Abs were found to bind autologous SHIV_1157ipd3N4_ gp120, both heavy and light chains of the Abs were cloned into the pCDNA3.1+ expression vector for large scale expression. The complete recombinant IgG proteins were expressed by co-transfection of paired heavy and light chain expressing plasmids into Expi293F (Thermo Fisher, Waltham, MA) according to the protocol for the ExpiFectamine^TM^ 293 Transfection Kit. Following 4-6 days incubation, culture supernatants were filtered and recombinant Abs were purified with Protein A Agarose (Thermo Pierce, Waltham, MA). Abs were eluted with IgG Elution Buffer (Thermo Pierce, Waltham, MA) and collected in 1M Tris pH 8 solution. The antibody solution was exchanged against PBS and the purified Abs were stored at -80°C.

#### Site-directed mutagenesis

Mutations in the SHIV_1157ipd3N4_ *env* gene were introduced by Site-Directed mutagenesis PCR (Gao et al., 2020). Briefly, a pair of oligonucleotide primers containing the desired mutation were used for PCR amplification using KOD-plus DNA polymerase (TOYOBO, Osaka). After amplification, the parental plasmid DNA template in the mixture was digested by Dpn I (Takara, Kusatsu, Shiga) treatment and the products with mutations were remained. Finally, the PCR products were transformed into DH5α competent cells and all *env* mutant plasmids were confirmed by sequencing.

#### Generation and titration of HIV-1 Env pseudoviruses

The Env pseudoviruses were generated by co-transfecting 8 μg of the *env* expression plasmid and 16 μg of the *env*-deficient HIV-1 backbone (pSG3ΔEnv) in to 293T cells using Lipofectamine 2000 transfection reagent (Invitrogen, Grand Island, NY) in T75 flask according to the manufacturer’s protocol. Cell culture supernatants were harvested 48-72 h following transfection and the subpackaged virus stock were stored at −80°C. The pseudoviruses were titrated by 11 5-fold serial dilution of each virus stock used to infect 1 × 10^4^ TZM-bl cells DMEM containing 10 µg/mL of DEAE-dextran, and a dilution which gave a relative luminescence units (RLU) signal that around 100 fold above average cell only wells was chosen to apply in neutralizing assay.

#### Neutralizing Assay

Neutralization activities of the antibodies were detected on TZM-bl cells as previously described (Gao et al., 2018). Briefly, purified antibodies (50 µg/ml) were serially diluted at a 1:3 ratio with DMEM. After incubation with certain concentration of Env-pseudoviruses at 37°C for 1 h, the 1 × 10^4^ TZM-bl cells in DMEM containing 10 µg/mL of DEAE-dextran were added into each well to be infected. After 48-72 h incubation, 200 µL of supernatant was removed and 40 µL of luciferase substrate was added into each well to test the RLU. The 50% inhibitory concentrations (IC_50_) were defined as antibodies concentration at which RLU were reduced by 50% compared with average RLU of virus control after subtraction of background RLU of cell control. A response was considered positive for neutralization if the IC_50_ concentration was lower than 50 µg/mL. The further neutralizing breadth and potency of J038 was evaluated on a panel of 208 circulating HIV-1 strains with microneutralization assay as previously described performed with an automated workstation (Doria-Rose et al., 2013; Sarzotti-Kelsoe et al., 2014) and the IC_50_ /IC_80_ was determined.

#### Expression and purification of HIV-1 Env proteins

The gp120 monomer and BG505 UFO trimer (Kong et al., 2016) were produced by transient transfection of plasmids into 293-6E suspension cells (NRC, Ottawa, ON) in serum-free medium, using a high-density transfection protocol. Briefly, 293-6E cells were thawed and incubated with OPM-293 CD05 Medium (OPM Biosciences, Shanghai) in a shaker incubator at 37 °C, with 100 rpm. and 5% CO_2_. The transfection procedure has been previously described (Sellhorn et al., 2009). Briefly, 0.25 mg of Env plasmid in 2.5 mL of OPM-293 CD05 Medium was mixed with 1 mL of PEI (1.0 mg/mL) in 2.5 mL of OPM-293 CD05 Medium. After incubation for 15 min, the DNA-PEI complex was directly added to 200 mL 293-6E cells. Culture supernatants were collected 3 days after transfection, then clarified by centrifugation at 3,000 rpm for 1 h and filtered using 0.45-μm filters (Thermo Scientific, Waltham, MA). The Env proteins were extracted from the supernatants using Lentil Lectin Sepharose^TM^ 4B (GE Healthcare, Boston, MA). The bound proteins were eluted with PBS containing 500 mM methyl-α-D-mannopyranoside and then concentrated for use. The affinity-purified Env proteins were further purified by size exclusion chromatography (SEC) using a Superdex 200 Increase 10/300 GL column (GE Healthcare, Boston, MA).

The HIV-1 C1080 Env for structural studies was expressed by transient transfection in 293 Freestyle cells. Briefly, 1 mg of DNA was transfected into 1L of cells using Turbo293 transfection reagent, and the cells were allowed to grow at 37°C for 6 days. Following expression, the supernatant was cleared by centrifugation and filtration, Supernatant was passed through a protein A column preloaded with CD4-binding site antibody 3BNC117, which has been engineered to have an HRV3C cleavage site at the hinge region. The resin was washed with PBS until the absorbance at 280 nm reached 0 and resuspended with 5-10 ml PBS. 200 µl of HRV-3C protease was then added to the resin mixture and incubated at 4 °C overnight to release the Env-3BNC117 Fab complex. Column flow-through and PBS wash (3 column volumes) were collected, concentrated and further purified with a Superdex 200 Increase 10/300 column to remove excess 3BNC117 Fab and HRV3C protease.

#### Enzyme-linked immunosorbent assay

Briefly, the autologous gp120 monomer or BG505 UFO trimer was coated into plates with 200 ng/well overnight at 4°C. Plates were washed three times with PBST (0.1% Tween 20 in PBS) and then blocked by PBS with 3% BSA for 2 h at 37°C. Then 5-fold serially diluted antibodies from 50 μg/mL were added into the wells with 1% BSA in PBS for 1 h incubation at 37°C. Plates were washed 3 times with PBST and horseradish peroxidase (HRP) rabbit anti-monkey IgG antibody (Bioss, Beijing) were added at 1: 1000 dilution for 1 h at 37°C. 100 μL/well tetramethylbenzidine (TMB) substrate were added and incubated for 15 min at room temperature in the dark and the reaction was stopped by adding 50 uL of 2 M H_2_SO_4_. The readout was detected at a wavelength of 450 nm.

#### Biolayer interferometry (BLI) measurements

BLI experiments were performed using the OCTET Red96 system (FORTÉBIO, San Jose, CA ) to determine binding curves of monoclonal antibodies for BG505 UFO and autologous gp120 (Sok et al., 2014). Briefly, 10 ng/mL monoclonal antibodies were immobilized onto Anti-hIgG-Fc Capture (AHC) biosensors (FORTÉBIO, San Jose, CA ) . Six serial dilutions from 800 nM to 25 nM of each antigen were prepared and flowed as analyte in solution. The binding experiment was performed at 30°C using the following protocol: baseline 1 (300 s), load antibodies (300 s), baseline 2 (60 s), Ab-antigen association (300 s) and dissociation (300 s). Binding affinity constants were determined using OCTET software Data Analysis HT 9.0 (FORTÉBIO, San Jose, CA ).

#### Inference of unmutated and intermediate evolution antibodies

The clonal tree, ancestral intermediate sequences, and unmutated common ancestor (UCA) sequence for the JO31 clone were inferred using Cloanalyst paired heavy and light chain UCA inference and clonal tree reconstruction (Kepler, 2013).

#### Preparation of the antibody Fab fragment and antibody-Env complex

For production of the Fab fragments of J033 and J038, a HRV3C protease cleavage site was inserted at the hinge region of IgG heavy chain gene as previously described (Shen et al., 2020). Cell culture supernatants of J033 and J038 were harvested, filtered, and loaded on Protein A columns (GE Health Science). Antibodies were eluted with IgG elution buffer (Pierce) and immediately neutralized by addition of one tenth volume of 1M Tris-HCl pH 8.0. To generate Fab fragments, purified IgGs were digested with HRV3C overnight at 4°C. The mixtures were passed through protein A columns again to remove Fc fragment or undigested IgG and the flow-though, which contained antibody Fab, were collected and concentrated for preparation of antibody-Env complex.

To obtain the J038 or J033 Fab-Env complexes, purified Fab was mixed with the C1080-3BNC117 complex at a 3.6:1 molar ratio and passed through a Superdex 200 Increase 10/300 column to separate excess Fab and the J038- or J033-C1080-3BNC117 complex.

#### Cryo-EM grid preparation and data collection

A sample of the J038-Env-3BNC117 or J033-Env-3BNC117 complexes at 1 mg/ml was diluted 1 mg/ml, 0.5 mg/ml, and 0.25 mg/ml in PBS buffer. 3 µl of each sample applied to a glow discharged Quantifoil R 1.2/1.3 Cu 200 mesh grid (Glow discharger settings: 0.30 mbar, 15 mAMP, and 5 seconds). Grids were then plunge frozen in liquid ethane with an FEI vitrobot and stored in liquid nitrogen (Vitrobot settings: 4 °C temperature, 100 % humidity, blot time 2 seconds, blot force 0, wait time 5 seconds, blot total 1, drain time 0).

#### Cryo-EM data processing and refinement

Images of the J033-Env-3BNC117 or J038-Env-3BNC117 complexes were collected on Titan Krios at magnification 18k with super-resolution mode. The original sampling is 0.68Å per pixel. We used bin=2 when processing the data and the sampling is 1.36Å per pixel.

The workflow on both data sets is shown in Figure Sxx. Relion 3.08 was employed to process the data. The images were further downscaled for the preprocessing process: manual particle picking, 2D classification to generate 2D templates for later auto particle picking, and 3D classification to generate a low-resolution model. Then the templates were applied to original data to do auto particle picking. After following the workflow of Relion, we got 3.4Å structure without using CTF Refinement and Bayesian Polishing to further push the resolution. Correlations between cryo-EM maps and atomic models were assessed using phenix.mtriage (Afonine et al., 2018). UCSF Chimera was used for docking and visualization (Pettersen et al., 2004). The coordinates of HIV-1 BG505 SOSIP in complex with 3BNC117 complex (PDB 5V8L) was used as initial models for fitting the cryo-EM map. The initial model for J038 Fab variable region was obtained using the SAbPred server (Shen et al., 2020). The J033 Fab was modelled based on refined J038 coordinates. Iterative model building and real space refinement were carried out using Coot (Emsley and Cowtan, 2004) and Phenix to fit the coordinates to the electron density map. Molprobity (Davis et al., 2004) was used to validate geometry and check structure quality.

### QUANTIFICATION AND STATISTICAL ANALYSIS

All inhibitory values (IC_50_ and IC_80_) of antibodies were calculated with a formally validated Excel-based macro (Piehler et al., 2011). The phylogenetic tree of neutralization results for 208-panel was constructed with Clustal X2. The ELISA binding curves were fitted using GraphPad Prism 8. The affinity constants of BLI were determined using OCTET software Data Analysis HT 9.0. The sequence alignments of antibodies or env coding regions were processed with Seaview 5.0.1. Cryo-EM data were processed and analyzed using cryoSPARC and Relion. Cryo-EM structural statistics analysis were carried out using Phenix, Molprobity, EMringer, PDBePISA and Chimera.

### DATA AND CODE AVAILABILITY

The sequences of 44 pairs of isolated antibodies and inferred intermediates have been deposited in Genbank, the accession numbers are: MZ234477-MZ234576. Cryo-EM map and coordinates of the J033-HIV-1 Env complex has been deposited under the accession codes EMDB:EMD-24128, PDB:7N28; Cryo-EM map and coordinates of the J038-HIV-1 Env complex has been deposited under the accession codes EMDB:EMD-24071, PDB:7MXD.

