## Supplementary figures for "Augmenting Neutralization breadth against Diverse HIV-1 by increasing the Ab-Ag interface on V2"

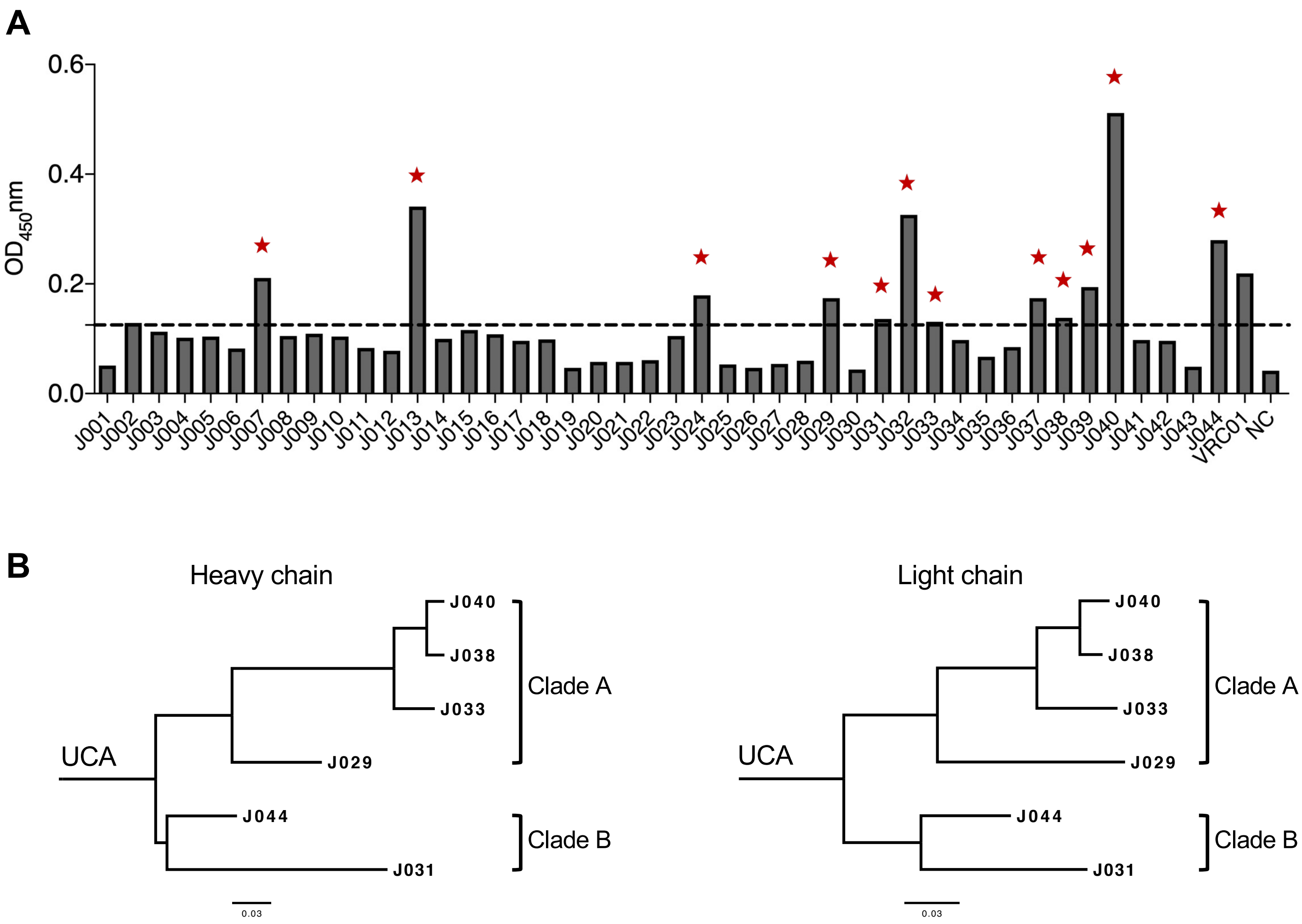


**Figure S1: Binding of newly isolated mAbs to autologous gp120 and Maximum-likelihood phylogenetic tree of the J038 lineage antibodies. Related to Figure 1.**

(A) Supernatants from the 293T cells transfected with paired antibody heavy/light chain linear DNA fragments (mAb ID indicated on the x-axis) were assayed by ELISA for binding to autologous Env (gp120). The mAbs that bind to gp120 are indicated by red stars. Supernatant from 293T cells transfected with VRC01 and from mock-transfection were used as positive control and negative controls (NC). The cutoff (2.1-fold of the NC value) for positive binding is indicated by the dash line.

(B) Maximum-likelihood phylogenetic tree of the heavy and light chain sequences of the J038 lineage antibodies. Unmutated common ancestor (UCA) of the six broadly neutralizing antibodies from the same gene family was inferred.

**Table S1. Lineage analysis and sequence characteristics of 12 monoclonal antibodies isolated from G1015R. Related to Figure 1.**

| | Heavy chain | | | | | Light chain | | | |
|---|---|---|---|---|---|---|---|---|---|
| mAb ID | IGHV | IGHD | IGHJ | CDR3 (aa) | SHM (%) | IGKV/LV | IGKJ/LJ | CDR3 (aa) | SHM (%) |
| J029 | 4-j*02 | 3-9*01 | 5-1*01 | AREVAEDDFGYYQPYYDS | 18 | K1-q*02 | K4*01 | QQYKALPLT | 20 |
| J031 | 4-j*02 | 3-9*01 | 4*01 | AREVPVDDYGYYLPYFDP | 22 | K1-b*06 | K4*01 | QQYVSMPLT | 17 |
| J033 | 4-j*02 | 3-9*01 | 4*01 | AGETPEDDFGYYQPYFKS | 23 | K1-q*02 | K4*01 | QHYKRLPLT | 20 |
| J038 | 4-j*02 | 3-9*01 | 4*01 | AGETPEDDFGYYQPYFKT | 22 | K1-q*02 | K4*01 | QHYRRLPLT | 16 |
| J040 | 4-j*02 | 3-9*01 | 4*01 | AGETPEDDFGYYQPYFKS | 23 | K1-q*02 | K4*01 | QHYKRLPLT | 15 |
| J044 | 4-j*02 | 3-9*01 | 4*01 | AREVAVDEYNYYAPYFDS | 11 | K1-b*06 | K4*01 | QQYKSLPLT | 14 |
| J039 | 4-n*01 | 4-4*02 | 4*01 | ATQSPLDGMSFGLNVA | 11 | K3-e*05 | K1*01 | HQYSDSVPWT | 11 |
| J024 | 4-e*01 | 5-5*02 | 5-1*01 | ARRRGDWLLSTKRTWFDV | 13 | K1-q*04 | K1*01 | QQGYNYPRT | 9 |
| J013 | 3-y*04 | 2-25*01 | 5-1*01 | IVGTKHCGDLYCPVAWFDV | 7 | K3-f*02 | K2*01 | QKQRDSPYT | 11 |
| J037 | 3-r*03 | 3-26*02 | 5-1*01 | VRGSHRGDYNRFFRGPKTSFDL | 19 | K1-r*01 | K2*01 | QHYYRSPYS | 6 |
| J032 | 3-r*03 | 3-21*01 | 5-1*01 | VRASHRGNYDRFFFSQKTWFDV | 10 | K3-f*02 | K2*01 | QEYDSYPYS | 6 |
| J007 | 3-r*03 | 3-26*02 | 5-1*01 | VRGSHRGDYDRFFRSPKTSFDL | 16 | L8-a*01 | L6*01 | TLYMGSGISM | 2 |

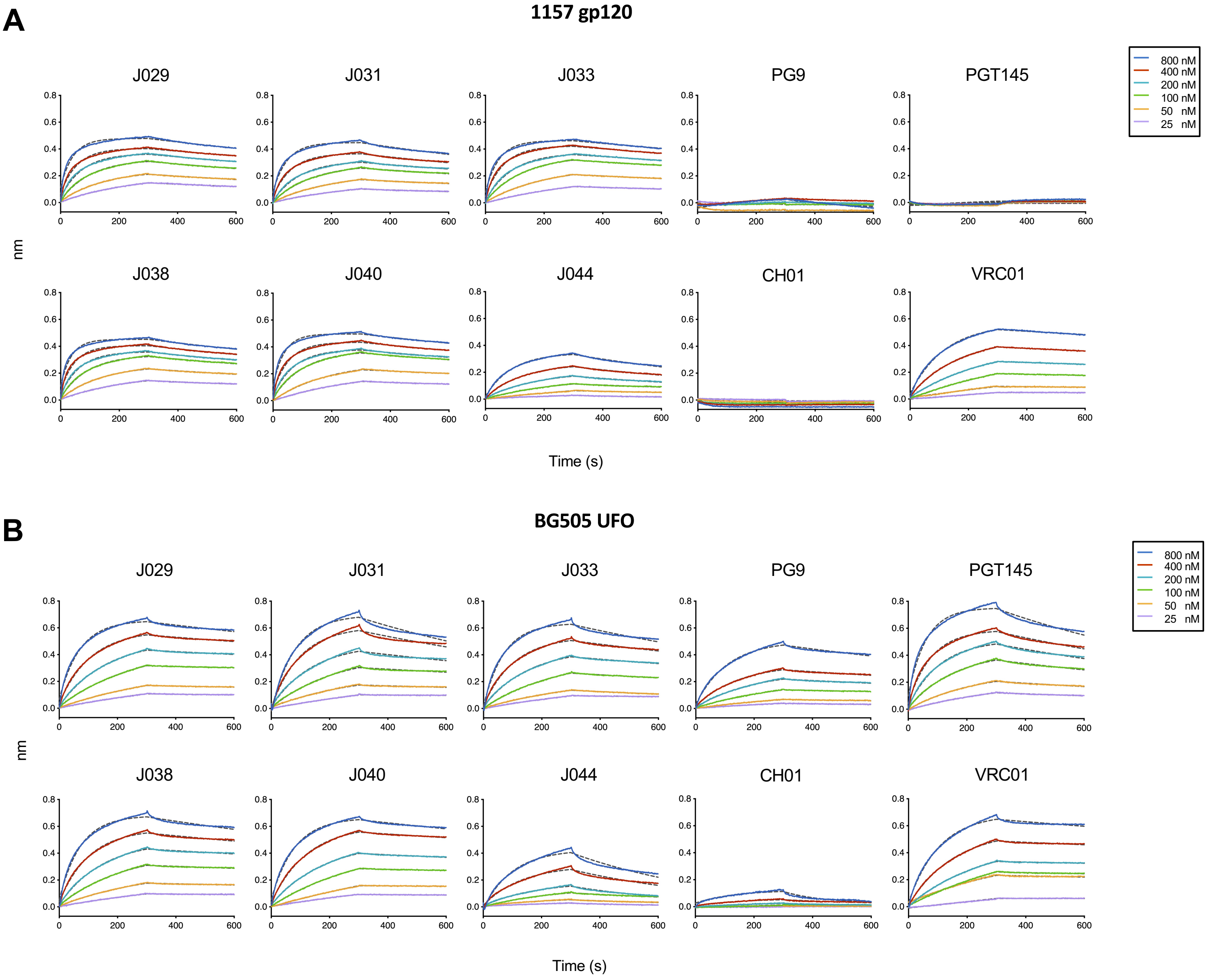


**C**

| Antigen | | J029 | J031 | J033 | J038 | J040 | J044 | PG9 | PGT145 | CH01 | VRC01 |
|---|---|---|---|---|---|---|---|---|---|---|---|
| | Kon (1/Ms) | 6.57E+04 | 4.80E+04 | 4.34E+04 | 7.95E+04 | 6.75E+04 | 1.55E+04 | N/A | N/A | N/A | 1.29E+04 |
| **1157 gp120** | Kdis (1/s) | 5.46E-04 | 4.57E-04 | 6.92E-04 | 6.01E-04 | 4.92E-04 | 1.08E-03 | N/A | N/A | N/A | 2.77E-04 |
| | KD (nM) | 8.3 | 9.5 | 15.9 | 7.6 | 7.3 | 69.5 | N/A | N/A | N/A | 21.4 |
| | Kon (1/Ms) | 2.31E+04 | 1.72E+04 | 2.04E+04 | 2.26E+04 | 1.96E+04 | 1.01E+04 | 1.32E+04 | 3.01E+04 | 1.31E+04 | 1.35E+04 |
| **BG505 trimer** | Kdis (1/s) | 3.45E-04 | 7.57E-04 | 6.76E-04 | 4.41E-04 | 3.06E-04 | 2.03E-03 | 6.06E-04 | 9.33E-04 | 3.44E-03 | 2.73E-04 |
| | KD (nM) | 15.0 | 44.0 | 33.1 | 19.5 | 15.6 | 201.0 | 45.8 | 31.0 | 263.0 | 20.2 |

**Figure S2: Binding affinity of the J038 lineage antibodies. Related to Figure 1.**

(A) The binding kinetics of six J038 lineage antibodies to autologous $SHIV_{1157ipd3N4}$ monomer gp120 were determined by BLI. The colored lines are the experimental curves for association and dissociation of the binding events, and the fitting curves were shown with gray dashed lines.

(B) The binding kinetics of six J038 lineage antibodies to autologous BG505 trimer were determined by BLI. The colored lines are the experimental curves for association and dissociation of the binding events, and the fitting curves were shown with gray dashed lines.

(C) Affinity constants for the J038 lineage antibodies. N/A; not applicable.

**Table S2. Neutralization activity of J038 against a panel of 208 viruses. Related to Figure 1.**

| Virus | Clade | $IC_{50}$ (μg/ml) | $IC_{80}$ (μg/ml) |
|---|---|---|---|
| 0260.v5.c36 | A | >50 | >50 |
| 0330.v4.c3 | A | 4.25 | 16.2 |
| 0439.v5.c1 | A | >50 | >50 |
| 3365.v2.c20 | A | 2.53 | 10.1 |
| 3415.v1.c1 | A | >50 | >50 |
| 3718.v3.c11 | A | 12.6 | 34.1 |
| 398-F1.F6.20 | A | 8.24 | 48.5 |
| BB201.B42 | A | 1.54 | 6.09 |
| BB539.2B13 | A | 6.00 | 22.4 |
| BG505.W6M.C2 | A | 8.67 | 29.5 |
| BI369.9A | A | 13.4 | 45.2 |
| BS208.B1 | A | 0.833 | 4.90 |
| KER2008.12 | A | 1.01 | 2.99 |
| KER2018.11 | A | 1.71 | 4.21 |
| KNH1209.18 | A | >50 | >50 |
| MB201.A1 | A | >50 | >50 |
| MB539.2B7 | A | 27.3 | >50 |
| MI369.A5 | A | >50 | >50 |
| MS208.A1 | A | 38.3 | >50 |
| Q23.17 | A | 9.06 | >50 |
| Q259.17 | A | 11.4 | >50 |
| Q769.d22 | A | >50 | >50 |
| Q769.h5 | A | >50 | >50 |
| Q842.d12 | A | 6.53 | 30.2 |
| QH209.14M.A2 | A | >50 | >50 |
| RW020.2 | A | >50 | >50 |
| UG037.8 | A | 3.25 | 8.33 |
| 246-F3.C10.2 | AC | 30.3 | >50 |
| 3301.v1.c24 | AC | 7.35 | 35.6 |
| 3589.v1.c4 | AC | 2.39 | 9.50 |
| 6540.v4.c1 | AC | 5.51 | 16.2 |
| 6545.v4.c1 | AC | 5.05 | 15.6 |
| 0815.v3.c3 | ACD | >50 | >50 |
| 6095.v1.c10 | ACD | 1.23 | 39.2 |
| 3468.v1.c12 | AD | >50 | >50 |
| Q168.a2 | AD | 12.6 | 34.7 |
| Q461.e2 | AD | >50 | >50 |
| 620345.c1 | AE | >50 | >50 |
| BJOX009000.02.4 | AE | 35.0 | >50 |
| BJOX010000.06.2 | AE | 25.1 | >50 |
| BJOX025000.01.1 | AE | >50 | >50 |
| BJOX028000.10.3 | AE | >50 | >50 |
| C1080.c3 | AE | 0.095 | 0.305 |
| C2101.c1 | AE | 4.27 | 11.5 |
| C3347.c11 | AE | >50 | >50 |
| C4118.09 | AE | 1.81 | 4.71 |
| CM244.ec1 | AE | 1.07 | 2.99 |
| CNE3 | AE | >50 | >50 |
| CNE5 | AE | 1.73 | 4.45 |
| CNE55 | AE | 14.2 | >50 |
| CNE56 | AE | >50 | >50 |
| CNE59 | AE | >50 | >50 |
| CNE8 | AE | >50 | >50 |
| M02138 | AE | >50 | >50 |
| R1166.c1 | AE | >50 | >50 |
| R2184.c4 | AE | >50 | >50 |
| R3265.c6 | AE | >50 | >50 |
| TH023.6 | AE | 0.009 | 0.067 |
| TH966.8 | AE | 1.36 | 4.59 |
| TH976.17 | AE | >50 | >50 |
| 235-47 | AG | >50 | >50 |
| 242-14 | AG | 6.23 | >50 |
| 263-8 | AG | 17.5 | >50 |
| 269-12 | AG | 49.9 | >50 |
| 271-11 | AG | >50 | >50 |
| 928-28 | AG | 6.48 | 42.1 |
| DJ263.8 | AG | 0.054 | 0.194 |
| T250-4 | AG | 0.839 | 2.79 |
| T251-18 | AG | >50 | >50 |
| T253-11 | AG | >50 | >50 |
| T255-34 | AG | >50 | >50 |
| T257-31 | AG | 11.5 | 28.6 |
| T266-60 | AG | 20.6 | >50 |
| T278-50 | AG | 12.3 | >50 |
| T280-5 | AG | 17.8 | 42.1 |
| T33-7 | AG | >50 | >50 |
| 3988.25 | B | 4.23 | 15.2 |
| 5768.04 | B | 14.2 | >50 |
| 6101.1 | B | >50 | >50 |
| 6535.3 | B | >50 | >50 |
| 7165.18 | B | 28.9 | >50 |
| 45_01dG5 | B | 7.04 | 27.4 |
| 89.6.DG | B | >50 | >50 |
| AC10.29 | B | 8.73 | 32.1 |
| ADA.DG | B | 0.347 | 1.64 |
| Bal.01 | B | 0.118 | 0.319 |
| BaL.26 | B | 0.163 | 0.545 |
| BG1168.01 | B | >50 | >50 |
| BL01.DG | B | >50 | >50 |
| BR07.DG | B | >50 | >50 |
| BX08.16 | B | 0.087 | 0.341 |
| CAAN.A2 | B | >50 | >50 |
| CNE10 | B | >50 | >50 |
| CNE12 | B | >50 | >50 |
| CNE14 | B | >50 | >50 |
| CNE4 | B | >50 | >50 |
| CNE57 | B | >50 | >50 |
| HO86.8 | B | 1.95 | 5.96 |
| HT593.1 | B | 3.79 | 15.0 |
| HXB2.DG | B | 0.010 | 0.032 |
| JRCSF.JB | B | 0.144 | 0.596 |
| JRFL.JB | B | >50 | >50 |
| MN.3 | B | 0.393 | 17.9 |
| PVO.04 | B | >50 | >50 |
| QH0515.01 | B | >50 | >50 |
| QH0692.42 | B | >50 | >50 |
| REJO.67 | B | 0.641 | 1.63 |
| RHPA.7 | B | 12.2 | >50 |
| SC422.8 | B | 13.8 | >50 |
| SF162.LS | B | >50 | >50 |
| SS1196.01 | B | 0.641 | 1.97 |
| THRO.18 | B | 3.82 | 16.6 |
| TRJO.58 | B | >50 | >50 |
| TRO.11 | B | 14.6 | >50 |
| WITO.33 | B | 0.334 | 1.58 |
| X2278.C2.B6 | B | 0.572 | 1.82 |
| YU2.DG | B | 8.99 | 24.5 |
| BJOX002000.03.2 | BC | 2.71 | 43.2 |
| CH038.12 | BC | >50 | >50 |
| CH070.1 | BC | 1.95 | 4.70 |
| CH117.4 | BC | 1.23 | 5.68 |
| CH119.10 | BC | 47.2 | >50 |
| CH181.12 | BC | >50 | >50 |
| CNE15 | BC | 5.83 | 22.2 |
| CNE19 | BC | 4.15 | 18.6 |
| CNE20 | BC | >50 | >50 |
| CNE21 | BC | >50 | >50 |
| CNE40 | BC | 0.231 | >50 |
| CNE7 | BC | >50 | >50 |
| 286.36 | C | 10.8 | 33.9 |
| 288.38 | C | 4.11 | 20.1 |
| 0013095-2.11 | C | 1.55 | 3.67 |
| 001428-2.42 | C | 2.02 | 4.93 |
| 0077.v1.c16 | C | 5.25 | 28.5 |
| 00836-2.5 | C | >50 | >50 |
| 0921.v2.c14 | C | 1.73 | 5.95 |
| 16055-2.3 | C | 2.25 | 5.89 |
| 16845-2.22 | C | >50 | >50 |
| 16936-2.21 | C | >50 | >50 |
| 25710-2.43 | C | 1.64 | 6.60 |
| 25711-2.4 | C | >50 | >50 |
| 25925-2.22 | C | >50 | >50 |
| 26191-2.48 | C | >50 | >50 |
| 3168.v4.c10 | C | 2.33 | 7.85 |
| 3637.v5.c3 | C | >50 | >50 |
| 3873.v1.c24 | C | 43.0 | >50 |
| 426c | C | >50 | >50 |
| 6322.v4.c1 | C | >50 | >50 |
| 6471.v1.c16 | C | >50 | >50 |
| 6631.v3.c10 | C | >50 | >50 |
| 6644.v2.c33 | C | 0.039 | 0.124 |
| 6785.v5.c14 | C | 1.01 | 3.50 |
| 6838.v1.c35 | C | 3.40 | 13.7 |
| 96ZM651.02 | C | >50 | >50 |
| BR025.9 | C | 0.480 | 3.30 |
| CAP210.E8 | C | 0.932 | 9.38 |
| CAP244.D3 | C | >50 | >50 |
| CAP256.206.C9 | C | 10.6 | 35.2 |
| CAP45.G3 | C | 0.259 | 1.02 |
| Ce1176.A3 | C | >50 | >50 |
| CE703010217.B6 | C | 1.54 | 4.66 |
| CNE30 | C | >50 | >50 |
| CNE31 | C | >50 | >50 |
| CNE53 | C | >50 | >50 |
| CNE58 | C | 2.66 | 10.6 |
| DU123.06 | C | 1.31 | 4.34 |
| DU151.02 | C | 2.03 | 6.05 |
| DU156.12 | C | 18.8 | >50 |
| DU172.17 | C | 13.2 | 44.0 |
| DU422.01 | C | >50 | >50 |
| MW965.26 | C | 0.094 | 0.790 |
| SO18.18 | C | 19.5 | >50 |
| TV1.29 | C | 0.889 | 3.86 |
| TZA125.17 | C | 30.7 | >50 |
| TZBD.02 | C | >50 | >50 |
| ZA012.29 | C | >50 | >50 |
| ZM106.9 | C | >50 | >50 |
| ZM109.4 | C | 1.00 | 7.06 |
| ZM135.10a | C | >50 | >50 |
| ZM176.66 | C | 0.677 | 2.46 |
| ZM197.7 | C | >50 | >50 |
| ZM214.15 | C | >50 | >50 |
| ZM215.8 | C | >50 | >50 |
| ZM233.6 | C | 0.096 | 0.459 |
| ZM249.1 | C | 22.8 | >50 |
| ZM53.12 | C | >50 | >50 |
| ZM55.28a | C | >50 | >50 |
| 3326.v4.c3 | CD | >50 | >50 |
| 3337.v2.c6 | CD | >50 | >50 |
| 3817.v2.c59 | CD | 12.5 | >50 |
| 191821.E6.1 | D | >50 | >50 |
| 231965.c01 | D | 14.6 | >50 |
| 247-23 | D | >50 | >50 |
| 3016.v5.c45 | D | >50 | >50 |
| 57128.vrc15 | D | >50 | >50 |
| 6405.v4.c34 | D | >50 | >50 |
| A03349M1.vrc4a | D | >50 | >50 |
| A07412M1.vrc12 | D | >50 | >50 |
| NKU3006.ec1 | D | >50 | >50 |
| UG021.16 | D | >50 | >50 |
| UG024.2 | D | 0.159 | 0.629 |
| P0402.c2.11 | G | >50 | >50 |
| P1981.C5.3 | G | 16.4 | >50 |
| X1193.c1 | G | 9.88 | 21.6 |
| X1254.c3 | G | >50 | >50 |
| X1632.S2.B10 | G | 12.0 | >50 |
| X2088.c9 | G | >50 | >50 |
| X2131.C1.B5 | G | 0.898 | 2.42 |
| SIVmac251.30.SG3 | NA | >50 | >50 |
| SVA.MLV | NA | >50 | >50 |

| | | | | | |
|---|---|---|---|---|---|
| <0.001 | 0.001-0.01 | 0.01-0.100 | 0.100-1.00 | 1.00-10.0 | >10.0 |

Neutralization potency was shown as $IC_{50}$ and $IC_{80}$ and color coded as indicated.

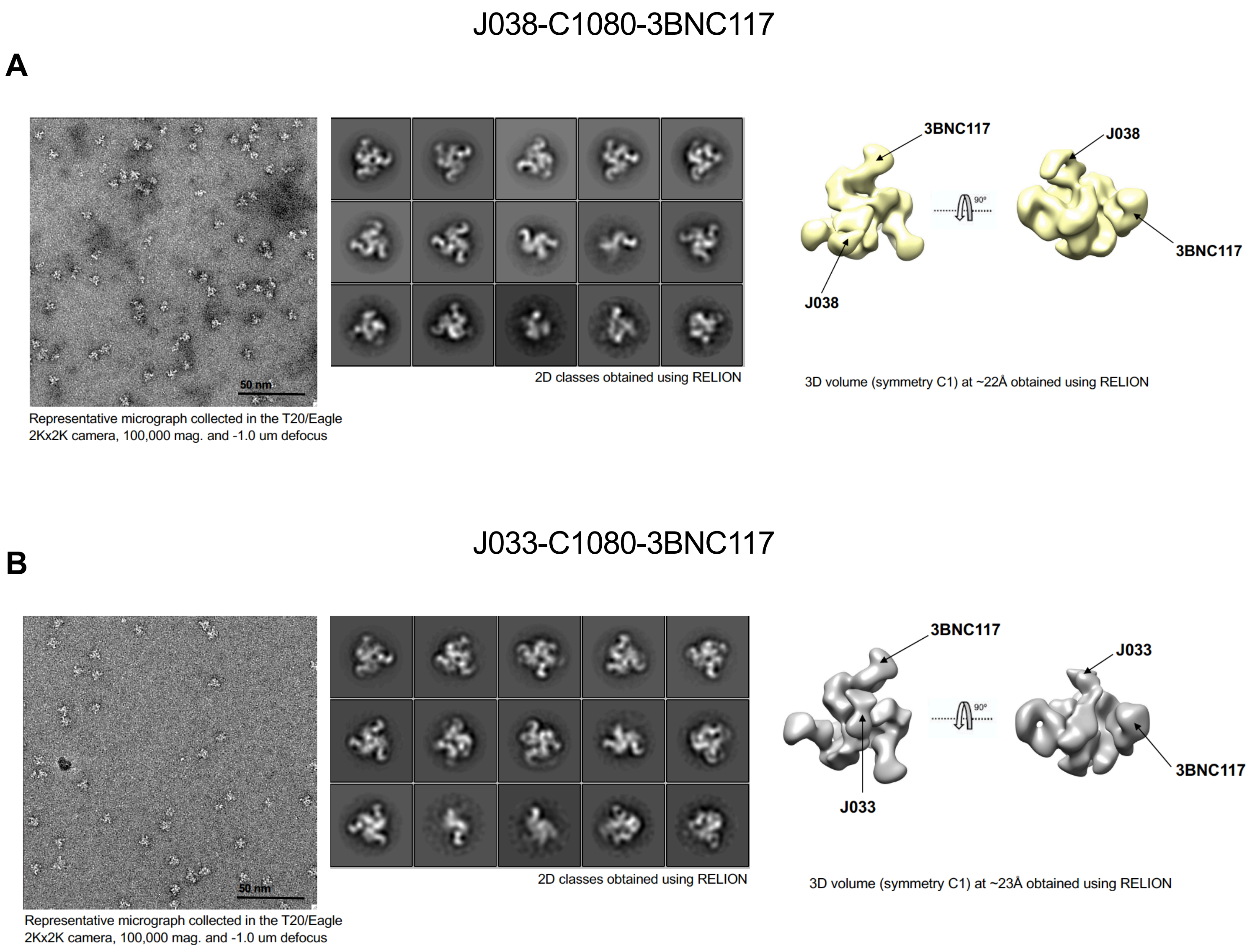


**Figure S3: Negative-stain EM and Cryo-EM of antibody-Env complex. Related to Figure 2.**
(A) Representative reference-free 2D class averages of J038-C1080 Env-3BNC117 complexes were shown at the middle panel.
(B) Representative reference-free 2D class averages of J033-C1080 Env-3BNC117 complexes were shown at the middle panel.

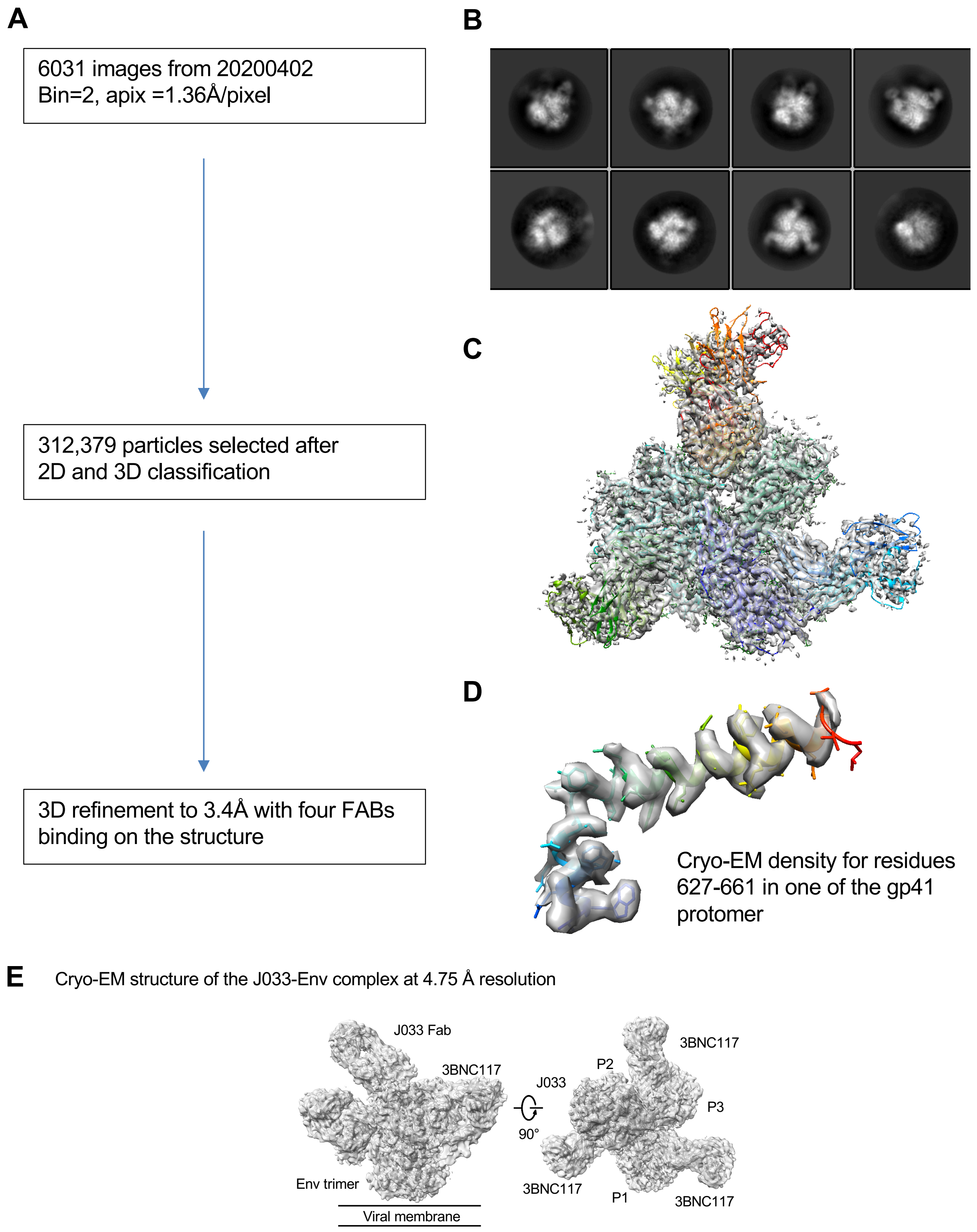


**Figure S4. Cryo-EM details of J038 in complex with HIV-1 Env. Related to Figure 2.**

A. Work follow of image processing.
B. 2D classes
C. Model fitting into the map, colored by chains
D. Representation region of the cryo-EM density with fitted residues
E. Cryo-EM structure of the Fab J033-Env complex, with EM reconstruction density shown in gray. The CD4-binding site antibody 3BNC117 was used to aid the resolution. Protomers of the trimer are labeled as P1, P2 and P3.

**Table S3. Cryo-EM data collection, refinement and validation statistics for J038 and J033 in complex with HIV-1 Env. Related to Figure 2**

| | C1080 Env in complex with J038 and 3BNC117 (EMDB:EMD-24071) (PDB:7MXD) | C1080 Env in complex with J033 and 3BNC117 (EMDB:EMD-24128) (PDB:7N28) |
|---|---|---|
| Data collection and processing | | |
| Magnification | 18,000 Super-res | 18000 Super-res |
| Voltage (kV) | 300 | 300 |
| Electron exposure ($e^{-}$/$Å^{2}$) | 50.0 | 50.0 |
| Defocus range (μm) | -1.6 to -3.0 | -1.6 to -3.0 |
| Pixel size (Å) | 0.68/1.36(bin=2) | 0.68/1.02 (bin=1.5) |
| Symmetry imposed | C1 | C1 |
| Final particle images (no.) | 312,379 | 383,584 |
| Map resolution (Å) | 3.40 | 4.20 |
| FSC threshold | 0.143 | 0.143 |
| Refinement | | |
| Initial model used (PDB code) | | |
| Model resolution (Å) | 3.64 | |
| FSC threshold | 0.143 | 0.143 |
| Map sharpening B factor ($Å^{2}$) | -89.5 | -129.8 |
| Model composition | | |
| Non-hydrogen atoms | 29496 | 29455 |
| Protein residues | 3519 | 3522 |
| Ligands | 145 | 140 |
| B factors ($Å^{2}$)(mean) | | |
| Protein | 186 | 88 |
| Ligand | 189 | 103 |
| R.m.s. deviations | | |
| Bond lengths (Å) | 0.003 | 0.003 |
| Bond angles (°) | 0.640 | 0.583 |
| Validation | | |
| MolProbity score | 1.73 | 1.97 |
| Clash score | 6.6 | 10.2 |
| Poor rotamers (%) | 0.58 | 0.03 |
| Ramachandran plot | | |
| Favored (%) | 94.7 | 93.0 |
| Allowed | 5.3 | 7.0 |
| Disallowed | 0 | 0 |

**Table S4. Interaction between between J038 and HIV-1 Env. Related to Figures 2, 3, 4 and 6**

A. Interface areas of paratope and epitope

| Antibody | HIV | Type | Epitope (Å$^2$) | | Paratope (Å$^2$) | |
|---|---|---|---|---|---|---|
| Heavy chain | Protomer P1 | Protein | 418 | | 409 | |
| | | Glycan 156 | | | | |
| | | NAG717 | 28 | | 25 | |
| | | NAG718 | 73 | | 70 | |
| | | Glycan 160 | | | | |
| | | NAG770 | 86 | | 75 | |
| | | NAG771 | 124 | | 99 | |
| | | BMA772 | 61 | | 62 | |
| | | MAN776 | 131 | | 99 | |
| | | MAN777 | 61 | | 56 | |
| | | MAN778 | 74 | | 71 | |
| Light chain | Protomer P1 | Protein | 201 | | 176 | |
| | | Glycan 156 | | | | |
| | | NAG718 | 17 | | 16 | |
| | Protomer P2 | Protein | 165 | | 140 | |
| Total surface area | | | 1439 | | 1298 | |
| | | by glycan | 655 | 45.5% | 573 | 44.1% |

B. Hydrogen bonds and salt bridges between J038 and HIV-1 Env

| | | Protomer 1 | Distance [Å] | Antibody |
|---|---|---|---|---|
| Heavy chain | 1 | F:LYS 171[ N ] | 3.72 | X:PHE 100A[ O ] |
| | 2 | F:LYS 171[ NZ ] | 3.74 | X:ASP 99[ O ] * |
| | 3 | F:TYR 173[ OH ] | 2.75 | X:ASP 100[ O ] |
| | 4 | F:ARG 166[ O ] | 3.53 | X:ARG 54[ NH2] |
| | 5 | F:ASP 167[ OD1] | 2.76 | X:TYR 33[ OH ] |
| | 6 | F:LYS 169[ O ] | 2.96 | X:TYR 100C[ N ] |
| | 7 | F:NAG 771[ O7 ] | 3.62 | X:TYR 100D[ OH ] |
| | 8 | F:NAG 771[ N2 ] | 3.34 | X:TYR 100D[ OH ] |
| | 9 | F:BMA 772[ O2 ] | 3.73 | X:ARG 54[ NH2] |
| | 10 | F:BMA 772[ O5 ] | 3.61 | X:ARG 54[ NH1] |
| | 11 | F:MAN 777[ O2 ] | 2.91 | X:ASN 30[ ND2] |
| Light chain | 12 | F:LYS 168[ NZ ] | 3.52 | Y:TYR 91[ O ] |
| | 13 | F:LYS 168[ NZ ] | 3.76 | Y:ARG 92[ O ] |
| | 14 | F:GLU 164[ OE1] | 3.88 | Y:ARG 93[ NH1] |
| | 15 | G:ASN 186[ ND2] | 2.58 | Y:ASP 1[ OD2] |
| | | Salt bridges | | |
| | | Protomer 1 | Distance [Å] | Antibody |
| Light chain | 1 | F:GLU 164[ OE1] | 3.88 | Y:ARG 93[ NH1] |

* UCA residues

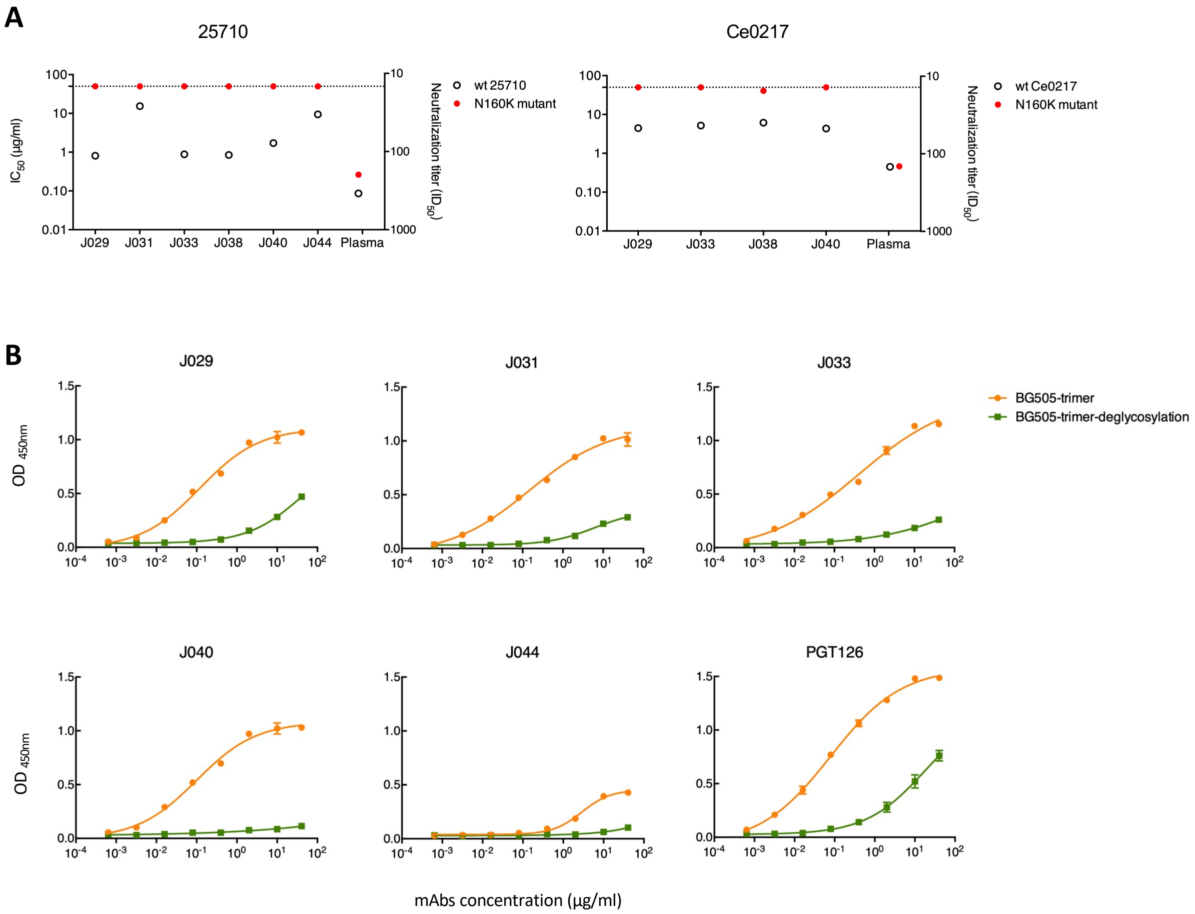


**Figure S5: The influence of glycan on antigen recognition of the J038 lineage antibodies. Related to Figure 4.**

(A) The N160K mutation rendered heterologous viruses 25710 and Ce0217 highly resistant to neutralization by the J038 lineage antibodies.

(B) The binding to the deglycosylated BG505 trimer by the J038 lineage antibodies was dramatically reduced. PGT126 serves as a positive control.

**A**

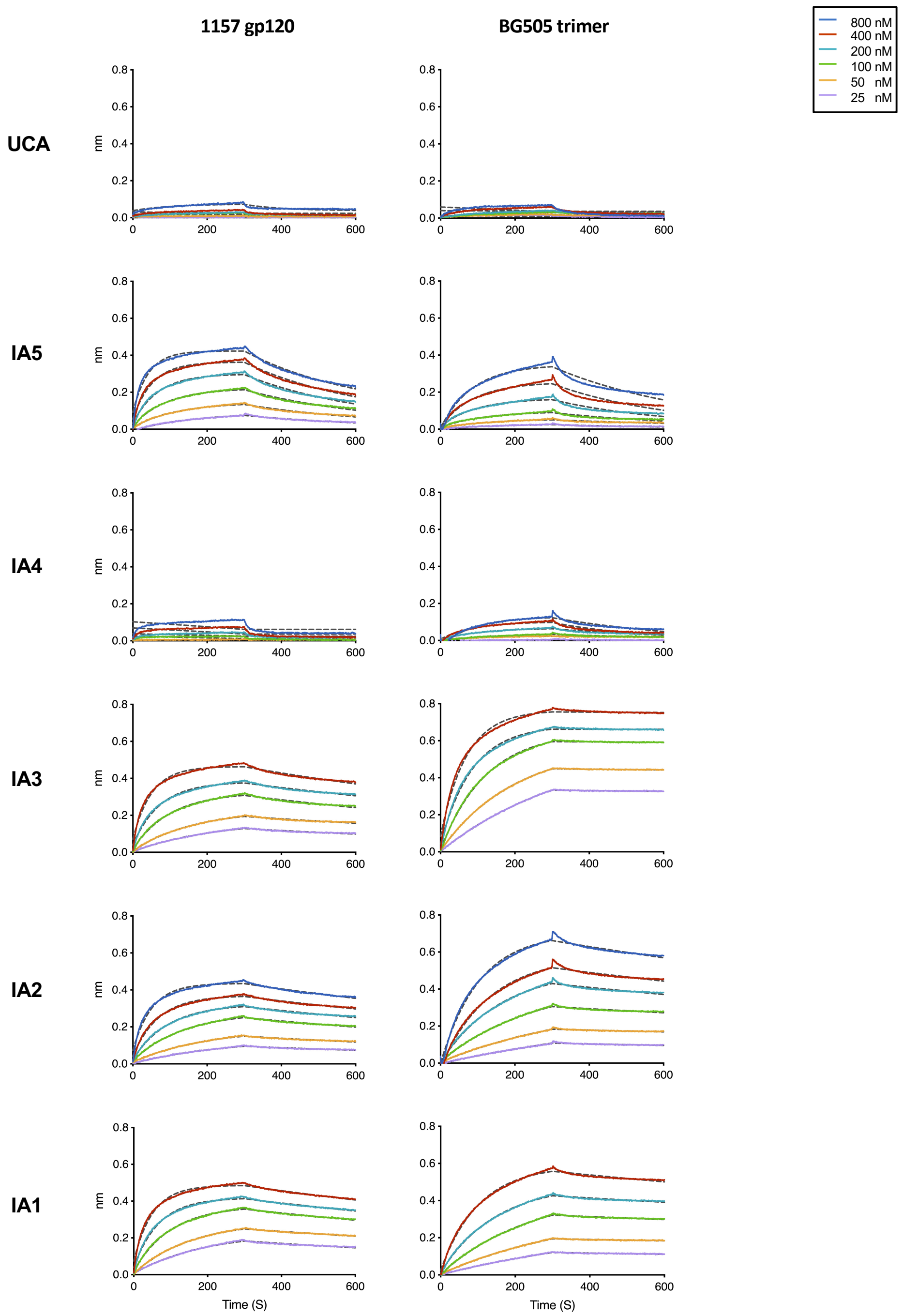


**B**

| Antigen | | UCA | IA5 | IA4 | IA3 | IA2 | IA1 |
|---|---|---|---|---|---|---|---|
| **1157 gp120** | Kon (1/Ms) | 6.09E+04 | 5.87E+04 | 1.38E+05 | 7.43E+04 | 4.18E+04 | 7.86E+04 |
| | Kdis (1/s) | 2.30E-03 | 2.41E-03 | 4.27E-03 | 7.42E-04 | 7.27E-04 | 6.17E-04 |
| | KD (nM) | 37.8 | 41.0 | 30.9 | 10.0 | 17.4 | 7.8 |
| **BG505 trimer** | Kon (1/Ms) | 3.66E+04 | 1.40E+04 | 7.92E+03 | 5.69E+04 | 1.25E+04 | 2.77E+04 |
| | Kdis (1/s) | 4.26E-03 | 2.84E-03 | 3.46E-03 | 6.17E-05 | 6.00E-04 | 3.43E-04 |
| | KD (nM) | 116.4 | 202.9 | 436.9 | 1.1 | 47.9 | 12.4 |

**Figure S6: Binding affinity of inferred UCA and IAs of the J038 lineage antibodies. Related to Figure 6**

(A) The binding kinetics of UCA and IAs of the J038 lineage antibodies to the autologous SHIV$_{1157ipd3N4}$ monomer gp120 or the BG505 trimer were determined by BLI. The colored lines are the experimental curves for association and dissociation of the binding events, and the fitting curves were shown with gray dashed lines.

(B) Affinity constants of the UCA and IAs of the J038 lineage antibodies.

### Table S5. Contributions of J038 and intermediate antibodies to binding of HIV-1 Env, related Figure 6

#### A. The interaction between J038 paratope residues and C1080 Env protein and glycans

| | | | | Buried surface area (Å2) | | | | | | | | | |
|---|---|---|---|---|---|---|---|---|---|---|---|---|---|
| | | | | Protein | Glycan N156 | | Glycan N160 | | | | | | |
| | Residue | Amino acid | Hydrogen bond and salt bridge | | glycan 717 | 718 | 770 | 771 | 772 | 776 | 777 | 778 | Total per residue |
| Heavy chain | 28 | X:ALA28 | | | | | | | | | 3.4 | | 3.4 |
| | 30 | X:ASN30 | | | | | | | 16.3 | 29.5 | 35.4 | | 81.2 |
| | 31 | X:ASP31 | | | | | | 16.6 | 4.5 | 1.2 | | | 22.3 |
| | 33 | X:TYR33 | H | 39.0 | | | | | | | | | 39.0 |
| | 50 | X:ARG50 | H | 36.3 | | | | | | | | | 36.3 |
| | 52 | X:SER52 | | 1.5 | | | | | | | | | 1.5 |
| | 54 | X:ARG54 | | 6.0 | | | | 27.3 | 41.6 | 35.1 | | 62.2 | 172.2 |
| | 55 | X:ASP55 | | 2.0 | | | | | | | | 7.0 | 9.0 |
| | 57 | X:TYR57 | | 33.9 | | | | | | | | | 33.9 |
| | 59 | X:GLU59 | | 16.6 | | | | | | | | | 16.6 |
| | 76 | X:TRP76 | | | | | | | | 33.0 | 17.2 | 2.2 | 52.4 |
| | 101 | X:PRO101 | | | | | | 7.5 | | | | | 7.5 |
| | 102 | X:GLU102 | | | | | | 5.1 | | | | | 5.1 |
| | 103 | X:ASP103 | H | 10.9 | | | | | | | | | 10.9 |
| | 104 | X:ASP104 | H | 38.4 | 25.5 | 25.1 | | | | | | | 89.0 |
| | 105 | X:PHE105 | H | 27.1 | | 45.1 | | | | | | | 72.2 |
| | 106 | X:GLY106 | | 23.4 | | | | | | | | | 23.4 |
| | 107 | X:TYR107 | H | 80.6 | | | 57.1 | 23.1 | | | | | 160.8 |
| | 108 | X:TYR108 | | 47.4 | | | 18.4 | 19.8 | | | | | 85.6 |
| | 109 | X:GLN109 | | 17.3 | | | | | | | | | 17.3 |
| | 110 | X:PRO110 | | 28.6 | | | | | | | | | 28.6 |
| Light chain | 1 | Y:ASP1 | H | 25.8 | | | | | | | | | 25.8 |
| | 26 | Y:THR26 | | 6.3 | | | | | | | | | 6.3 |
| | 27 | Y:GLN27 | | 42.3 | | | | | | | | | 42.3 |
| | 28 | Y:GLY28 | | 20.4 | | | | | | | | | 20.4 |
| | 32 | Y:ASP32 | | 8.74 | | | | | | | | | 8.7 |
| | 52 | Y:PHE52 | | | | 15.9 | | | | | | | 15.9 |
| | 91 | Y:TYR91 | H | 21.9 | | | | | | | | | 21.9 |
| | 92 | Y:ARG92 | H | 58.3 | | | | | | | | | 58.3 |
| | 93 | Y:ARG93 | H, S | 87.3 | | | | | | | | | 87.3 |
| | 94 | Y:LEU94 | | 36.9 | | | | | | | | | 36.9 |
| | 96 | Y:LEU96 | | 7.5 | | | | | | | | | 7.5 |

UCA residue

#### B. Increased binding surface areas to glycan by mutations accumulated on intermediate antibodies

| Paratope | Heavy chain (Å2) | Light chain (Å2) | Increasing interface (Å2) | |
|---|---|---|---|---|
| Contribution by UCA | 497.6 | 129.0 | 626.6 | 48% |
| By mutations in IA5 | 237.4 | 96.0 | 333.4 | 26% |
| By mutations in IA3 | 144.6 | 22.2 | 166.7 | 13% |
| By mutations in IA2 | 7.5 | 87.3 | 94.8 | 7% |
| By mutations in IA1 | 0 | 0 | 0 | 0% |
| Total | 968.2 | 331.3 | 1299.5 | |

```
Rhesus 4-j*02  QLQLQESGPGLVKPSETLSLTCAVSGGSISSNYWSWIRQPPGKGLEWIGRISGSGGSTDYNPSLKSRVTISTDTSKNQFSLKLSSVTAADTAVYYCARDL
Human  4-4*08  -V--------------------T--------Y-----------------Y-YT--.--N------------V----------------------------
```

**Figure S7: Comparison between the VH germline gene sequence (4-j*02) of the J038 lineage antibodies in rhesus macaques and its orthologous gene sequence (4-4*08) in humans. Related to Figure 6.**
